# On the evolution of sexual receptivity in female primates

**DOI:** 10.1101/727875

**Authors:** Kelly Rooker, Sergey Gavrilets

## Abstract

There has been much interest in the evolutionary forces responsible for, and underlying the diversity in, female primate reproductive cycles. Some primate species are like humans, sexually receptive to mating throughout their entire estrus cycle, while other species are the opposite, receptive for mere hours out of their several-week cycles. Why is there such prominent variation in sexual receptivity length among primate species? Here we examine the evolutionary trade-offs associated with sexual receptivity length using mathematical modeling. We investigate how various factors, including having ovulation signs present vs. concealed ovulation, female physiological costs, and group size, each influence the length of females’ receptive periods. We find that both continuous receptivity and very short lengths of receptivity are able to evolve. Our model predicts that increasing the impacts of infanticide will increase the length of the female receptive period, emphasizing the possible importance of paternity confusion. Similar effects can also be achieved by increasing the non-genetic benefits provided by males. Overall, our work offers a theoretical framework for understanding the evolution and diversity of mating traits in female primates.

## Introduction

Research on the unique behavioral changes associated with the estrus cycle of female mammals has been of great interest for over a century. As early as 1900, scientists were interested in when and why females were willing to receive male mates, noting that in most species there was a “special period of sexual desire…and only at that time, the female is willing to receive the male” (Heape, 1900, p.6).

### Definitions

In general, there are three main characteristics used to describe the changes through which female mammals go when they are in estrus: sexual attractivity, proceptivity, and receptivity (Beach, 1976). Although the focus of this paper is on female sexual receptivity, much of the work surrounding the estrus cycles of female mammals has instead been in relation to *sexual attractivity*, that is non-behavioral cues, such as sexual swellings, bright colorations, etc. around the female’s genital region, that are used to signal probable times of ovulation. Concealed ovulation, also of much interest, is the lack of any such visual signs around a female’s time of ovulation. *Proceptivity* refers to behavioral cues displayed by females in order to initiate and/or maintain sexual interactions with males. Finally, *receptivity* is the willingness of a female to accept a male and allow copulation/intravaginal ejaculation with him to occur (Dixson, 2012).

Hence, attractivity can be thought of as the value to a male of a female being a sexual stimulus, and typically gets measured for primates in the field via male behaviors: frequency of approaches by males to a female, frequency of attempts by males to mount a female, etc. (Hrdy and Whitten, 1987). Proceptivity instead gets measured via female behaviors: counts of affiliative behaviors (e.g., a female moving to sit near a male or to stay in his vicinity), more direct sexual solicitation (e.g., a female presenting her hindquarters to a male or making species-specific vocalizations, facial expressions, and/or gestures like lip-smacking in baboons or head-bobbing in rhesus monkeys), investigation of the male’s anogenital region, female grooming of the male, etc. (Beach, 1976). (See (Dixson, 2012, p.133) for a more complete list of proceptive behaviors in female primates.)

Receptivity is harder to measure in the field since it requires a male approaching and wanting to mate. However, it can be seen by the female shifting to a specific stance allowing for penile insertion or ejaculation (depending on the species), remaining stationary in that position, etc. (Hrdy and Whitten, 1987). Receptivity, even when the female is most fertile, is not a passive action. Females may avoid or actively refuse male mount attempts, or terminate the mount prior to ejaculation. In the field, receptivity in primates often gets measured by counting such refusals, acceptances, and terminations of male mounts by females (Dixson, 2012).

Also, it is important to note that even in species with long lengths of receptivity, matings will not necessarily be evenly distributed throughout such periods of receptivity. Things like female desire and female-initiated matings may increase around their time of fertility, despite a female being receptive for longer proportions of her cycle (Hrdy, 1997).

### Receptivity in Non-Primate Mammals

Receptivity as an aspect of primate behavior is much different from the vast majority of nonprimate mammals (Beach, 1976). In most non-primate mammals, receptivity is limited to the peri-ovulatory (i.e., around the time of ovulation) phase of the female’s estrus cycle, and strongly dependent on hormones, including estrogen and progesterone. Prosimian primates (lemurs, galagos, etc.) also tend to be more like non-primate mammals in that they only have limited periods of sexual receptivity during their peri-ovulatory phase (Dixson, 2012). Not only is this time of receptivity typically brief in non-primate mammals and prosimians, but in many species (including galagos), the vagina is actually fused shut, or otherwise completely covered, outside the female’s receptive period, making mating outside of this time physically impossible (Hrdy and Whitten, 1987).

In addition, in many non-primate mammals and prosimian primates, ovariectomy (surgical removal of the ovaries and hence also removal of the ovary-produced hormones estrogen and progesterone) results in a complete lack of sexual receptivity in the female, with females even attacking males attempting to copulate with them in the case of galagos. This is quite different from the anthropoid primates (including humans), in which copulations will still occur, even after removal of the ovaries (Dixson, 2012).

### Variation in Receptivity among Primates

Among all species of primates, there is great variability in the duration of the estrus cycle where the female is receptive. Some species are like humans or bonobos in which individual females are receptive throughout their entire cycle, while other species are like gorillas or the prosimians, in which individual females are receptive exclusively during the middle of their cycles around their time of ovulation (Beach, 1976). In some species, females will initiate the majority of all matings (e.g., in *Gorilla*, *Ateles*, *Alouatta*, *Cebus*, and *Rhinopithecus*), while in other species females initiate 0% of all mounts (e.g., in greater galagos and owl monkeys). Other primate species lie more in between, with females initiating 18% of all mounts in free-ranging chimpanzees, 50% in vervet monkeys, and 69% in lion-tailed macaques (Dixson, 2012).

### Other Aspects of Receptivity

Species whose females are receptive throughout their entire cycle are referred to as being ‘continuously receptive’. Many species engage in what is called ‘situation-dependent receptivity’, i.e., when a female becomes receptive despite not being in the middle of her cycle. This most frequently occurs when a female encounters an unfamiliar male. For example, non-receptive females (including in some cases even pregnant females) in gray langurs and gelada baboons will display both proceptive and receptive behaviors within days of new males invading their group (Hrdy and Whitten, 1987).

Longer receptivity is clearly *less* important in species where forced copulations occur (Clarke et al., 2009). However, in many primate species, the female takes an active role in mating via proceptive behaviors and many males wait for such behaviors prior to attempting to mate (Beach, 1976). On the other hand, longer receptivity could be *more* important in select species where ‘food-for-sex’ is the norm (i.e., the female receives provisionings or other resources from a male in exchange for her copulating with him) (Stockley and Bro-Jorgensen, 2011). Similarly - and more commonly - the female could instead be trading copulations with a male for protection of herself against predators, harassment, etc., which could also provide selection pressure on the lengthening of receptivity. Alternatively, females could increase their fitness by mating with males more likely to provide paternal care to their offspring (Clutton-Brock, 2016). Note all these different costs and benefits of receptivity can be thought of as encapsulated together in a single ‘benefit-to-cost’ ratio, what we term later the ‘benefit of paternal male quality’.

### Female Competition for *Preferred* Mates

As stated in a review paper on female-female competition, “Female competition for the sperm of preferred (or competitively successful) males could be a potentially widespread but previously overlooked evolutionary force” (Stockley and Bro-Jorgensen, 2011, p.361). Indeed, receptivity may function less in securing females *any* mate, but rather in securing them their *preferred* mate (i.e., a high-quality male). In these situations, females are trying to concentrate paternity of their offspring in that preferred mate. At least in some contexts, sperm limitation may play a role (Clutton-Brock, 2016). In both gelada baboons and hamadryas baboons, there is a negative relationship between conception rate and the ratio of estrus females to males. Similarly, among gorillas, females will receive fewer copulations when other females are in estrus simultaneously (Stockley and Bro-Jorgensen, 2011). Note that such preference in females for ‘high-quality males’ can be thought of not only as a preference for good genes, but also in maturity, rank, fertility, protection, investment, vocal or visual displays, etc. (Clutton-Brock, 2016). Surbeck et al. showed that, when comparing bonobos to chimpanzees, male bonobos have a higher reproductive skew and stronger relationship between dominance rank and reproductive success, despite female bonobos having the longer periods of receptivity (Surbeck et al., 2017).

In general, there exists a trade-off between the benefits of a female being receptive for a longer amount of time vs. a female being receptive for shorter. A female with a shorter period of receptivity would have less time in her cycle to mate, which helps both concentrate paternity in high-quality males and also avoid the physiological costs of having longer receptivity. On the other hand, a female with a longer period of receptivity would have more time to mate with multiple males in one cycle, helping to confuse paternity (van Schaik et al., 1999). Possible benefits of mating with multiple males in one cycle include reduced risk of infertility, increased protection of offspring against predators, increased investment of offspring by multiple males (but see (Soltis and McElreath, 2001)), and genetic diversity (Clutton-Brock, 2016).

### Infanticide

Another important possible benefit to females from both paternity concentration and paternity confusion - and hence also a possible effect on female receptivity length - comes from infanticide. Infanticide, the killing of a female’s offspring prior to weaning, is extremely detrimental to a female’s fitness (for example, see Hrdy (1979); Paul (2002); Pradhan and van Schaik (2008); Van Noordwijk and van Schaik (2000); van Schaik (2000); van Schaik et al. (2004); Zipple et al. (2017)). However, committing an act of infanticide may benefit any male who is not the father of that particular offspring, and hence infanticide is often seen when a new male takes over a group (Pradhan and van Schaik, 2008). Since many female primates have prolonged lactational amenorrhoea (van Schaik et al., 1999), a male killing a female’s offspring results in bringing that female back to fertility sooner by prematurely ending her lactation (Lovejoy, 2009; Palombit, 1999). The infanticidal male can then subsequently mate with this female, meaning his offspring will be able to be born sooner than would otherwise be possible had he never committed infanticide (Stallman and Froehlich, 2000). Although infanticidal events are typically rare, they have been documented in many primate species (Hrdy, 1979; Opie et al., 2013), and constitute up to 64% of all infant mortality in some primate species (van Schaik et al., 1999).

Indeed, since males’ decisions about whether to help or harm an infant are primarily based on their mating history, one prominent counterstrategy used by females to protect against infanticide is the manipulation of males’ probabilities of paternity, either actual or perceived. For example, if infanticide is likely to only be committed by one male in a group, it may be in females’ best interests to concentrate their future offspring’s paternity in him. This would help ensure that he believes he is the father of her offspring, and hence he would be much less likely to kill the offspring, and may even go out of his way to help protect it should any other male attempt to commit infanticide against that offspring (van Schaik et al., 1999). Observational and genetic data both confirm that infanticide is rarely committed by the father of that offspring (Clutton-Brock, 2016).

Conversely, paternity confusion could also play a role in protecting an offspring from infanticide. If infanticide may be committed by multiple males in the group, or any new outsider male who takes over the group, it would instead likely be in the female’s best interest to mate with as many of those males during one cycle as she possibly can, in order to help confuse paternity, i.e., make multiple males think they may be the father of her offspring (van Schaik et al., 1999). Indeed, infanticide could have provided the main selection pressure favoring situation-dependent receptivity, including continuous receptivity. Such a female with the capacity to facultatively and opportunistically solicit sexual behavior would be better equipped to manipulate males’ probable paternities, thus lowering the risk that her infant might be killed (Hrdy, 1979).

### Research Questions

In situations with no fossil record, as is the case with female receptivity, scientists must often rely upon comparative analyses and evolutionary modeling in order to help understand such phenomena. The application of the comparative approach to female primate sexual receptivity has, so far, been limited. Hrdy and Whitten (1987) collected much data on female primate receptivity, proceptivity, sexual swellings/colorations, male behaviors, breeding patterns, etc. However, their work was meant as more of an exposé of all the variation that is present among primate species rather than a true comparative analysis between different traits. van Schaik et al. (1999) were primarily concerned with infanticide and those sexual behaviors relating to defense against infanticide. Their collected data and statistical analyses focused on correlations among traits and behaviors present in today’s species, rather than on their evolutionary origins. Finally, (Stockley, 2002) looked only at the specific relationships between female receptivity length and male baculum length and penile spinosity in primates.

Given the complex evolutionary trade-offs between factors affecting female receptivity, theoretical insight based on evolutionary modeling is warranted. However, to our knowledge there has yet to be any mathematical modeling work done on the evolution of primate sexual receptivity. It is our aim to address this gap. In particular, we are interested in the following questions: How does length of sexual receptivity contribute to female reproductive success? What factors facilitate the evolution of continuous receptivity and/or very short lengths of receptivity? What role could infanticide have in the evolution of receptivity? How is receptivity linked to the evolution of sexual attractiveness, namely a female having obvious visual ovulation signs or concealed ovulation?

## Methods

Similar to Rooker and Gavrilets (2018), we construct an agent-based model using a population of individuals living in a large number of groups, each with *N* males and *N* females. See Appendix A for full details of the model. We assume generations to be discrete and non-overlapping. Females can differ genetically in both their visible ovulation signs present and their length of receptivity, while males differ in their quality to females, meaning any benefit from the male to a female and/or her offspring. We do not consider evolution in males, assuming instead that male traits are at a [stochastic] evolutionary equilibrium.

### Males

Variation in male quality is scaled by parameter *b*, which also characterizes female benefits. Such quality is explicitly split into a genetic component *y_g_* (GC) and a non-genetic component *y_ng_* (NGC). GC is due to the male’s genes being passed to the female’s offspring, while NGC is any non-genetic benefit to the female from mating with that male. For example, NGC could be protection provided to the female or increased food, provisioning, access to resources, etc. (as reviewed in detail in Clutton-Brock (2016)). We assume that *y_g_* correlates with each male’s rank (and hence his success in reproductive competition). We also allow for correlation *ρ* between *y_g_* and *y_ng_*. For example, *ρ <* 0 corresponds to the case when powerful males are less interested in providing any protection/provisioning to females. *ρ >* 0 instead corresponds to the case when being a more powerful male implies being better at protecting/provisioning females. Parameter 0 *≤ η ≤* 1 specifies the relative weight of NGC; with *η* = 0.5, male-provided genetic and non-genetic benefits are equally important for females.

### Females

We explicitly account for the female cycle, which we split into *D* discrete units of time. While such units could be hours, minutes, etc., we choose to call such units ‘days’ (for example, with 29 days in one female cycle). Each female is characterized by one genetically controlled trait, *r*, the length of time the female is receptive to mating (a non-negative, integer value). In addition, females may also be characterized by their visual ovulation signs, denoted *x*(*d*) for each day *d* of the cycle, thought of as overlapping curves. Visual ovulation signs are characterized by two evolvable, genetically controlled traits. Ovulation signs magnitude *m* is the maximum amount of ovulation signs a female has visible during her cycle (a non-negative, continuous value), while ovulation signs length *l* is the number of days a female has *some* amount of ovulation signs visible (a non-negative, integer value). We also account for the costs of having ovulation signs visible (scaled by parameter *c*) and the costs of receptivity (*c_r_*). For an example of what receptivity lengths *r* and ovulation signs *x*(*d*) could look like across a cycle, see Fig. 1.

For each female, the distribution of probabilities of fertilization across the cycle has the same shape but is randomly centered (meaning cycle synchrony may only occur probabilistically). Visual ovulation signs and receptivity are correlated with these days of fertility, but can each still evolve independently. This means we assume there exists at least some reliability of the ovulation signal (since a female’s peak day of fertility will also be her day of peak ovulation signs *and* her median day of receptivity), and that such reliability is the same in all females.

### Mating

For every unit of time, all males and all receptive females in each group enter the competition for mates. For computational simplicity, we assume each individual to mate exactly once on every unit of time (e.g., day). Males of higher rank (and GC) are more likely to mate with receptive females with stronger ovulation signs (higher *x*(*d*)), i.e., when the female is likely to be most fertile. Males of lower rank (and GC) mate with receptive females who have fewer ovulation signs visible, i.e., when the female is less likely to be most fertile. For empirical support of this assumption in primates, see Refs. Deschner et al. (2004); Domb and Pagel (2001); Higham et al. (2012). The strength of such assortment in our model is controlled by reproductive stochasticity parameters *∊_m_ ≥* 0 and *∊_f_ ≥* 0, in males and females respectively. With *∊_m_* = *∊_f_* = 0, mating pairs are formed deterministically so that the highest-GC male mates with the female with the most ovulation signs visible, and so on; this is the case of perfect assortment in mating. Increasing *∊_m_* increases stochasticity in the outcome of male competition for mating success, and thus reduces male reproductive inequality. Increasing *∊_f_* reflects a decrease in the reliability of the ovulation signal as well as a lack of opportunity or interest for fertile females to mate with a higher-GC male (e.g., as shown empirically in Amboseli baboons (Fitzpatrick et al., 2015)). With large *∊_m_*, *∊_f_*, mating becomes random.

**Figure 1:**
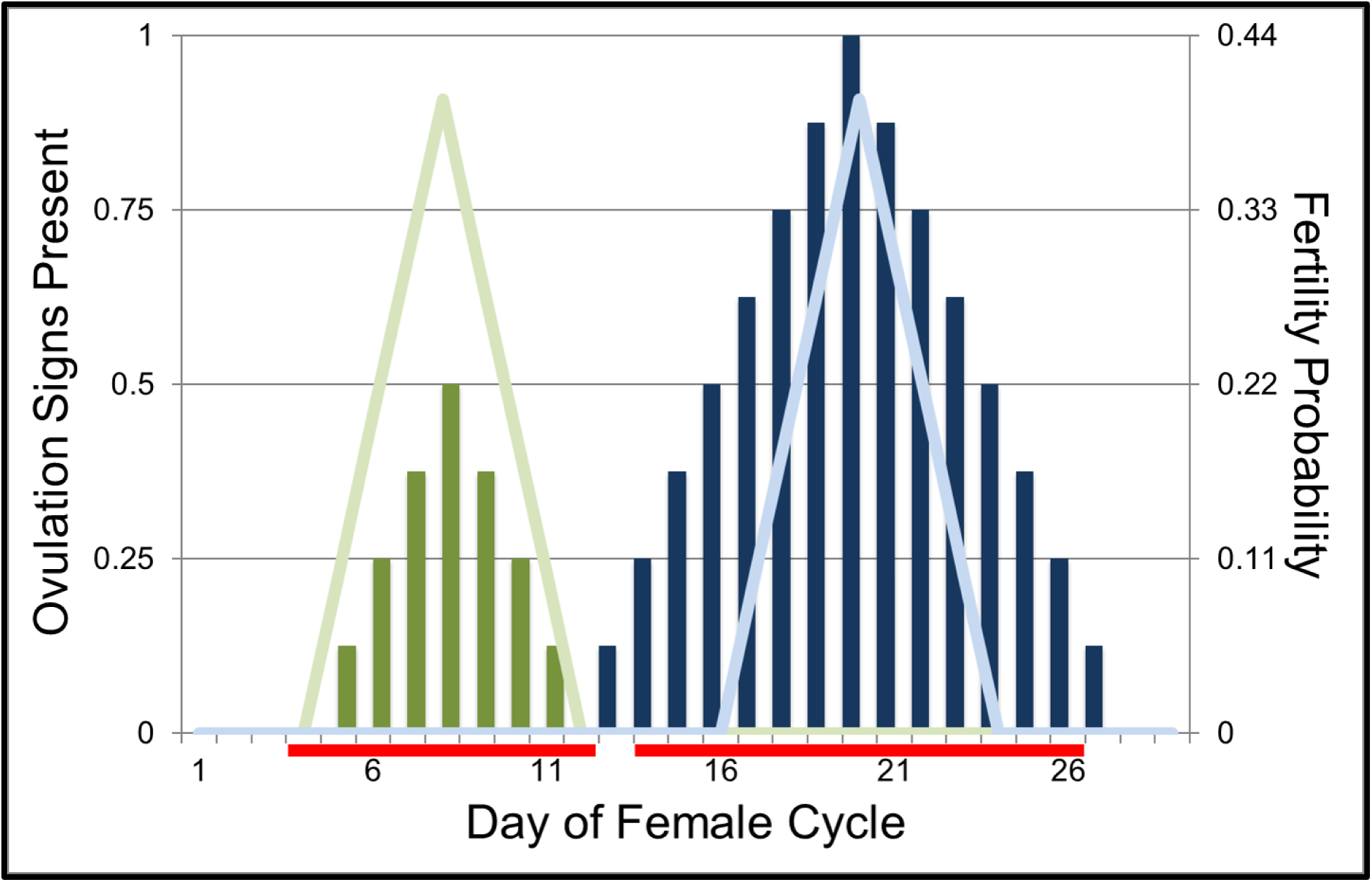
Example ovulation signs, fertility, and receptivity length for two females. Female 1 is denoted by green (*r* = 9, *m* = 0.5, *l* = 7), and Female 2 by blue (*r* = 13, *m* = 1, *l* = 15). Visible ovulation signs are depicted by the solid vertical bars, fertility probabilities by the lighter-colored lines, and receptivity by the red horizontal bars at the bottom of the graph. Ovulation is assumed to happen on Day 8 for Female 1 and Day 20 for Female 2. Note receptivity length and ovulation signs and magnitude can all vary independently between females, while fertility curves have the same shape for all females.

Mating is followed by offspring production and dispersal. For any female’s potential offspring, every male in the group will have an associated probability of paternity, ranging anywhere from zero to one. This quantity is determined by both the number of times that particular male mates with the female during her cycle, as well as her probability of fertilization on the days of each mating event.

### Infanticide

Modeling infanticide is similar to that in Rooker and Gavrilets (2018), which itself was adapted from van Schaik et al. (2004). Any male is able to help or harm (note ‘harm’ here could simply mean not helping) an offspring prior to the infant being weaned, although some males (e.g., those of higher quality, size, or strength) are able to do so more effectively. Our model does not distinguish between the threat of infanticide via an outsider male or a previously-subordinate male. Infanticide’s effects on offspring viability are scaled by parameters *α* (maximum benefit of protection from infanticide) and *β* (relative maximum cost of infanticide). An additional parameter 0 *≤ κ ≤* 1 determines the extent to which males take visible female ovulation signs into account when estimating their paternity in determining what actions to take regarding infanticide. *κ* = 0 means males estimate their probability of paternity solely by the number of matings with an offspring’s mother, while *κ* = 1 means males instead estimate paternity exclusively on the basis of the female’s visible ovulation signs during their time of mating.

### Female Fitness

Fitness is each female’s relative reproductive success, i.e., the expected number of offspring surviving to the age of reproduction, normalized to keep the total population size in each generation constant. Assuming additivity of cost and benefit effects for simplicity, we specify this female fitness (fertility) as

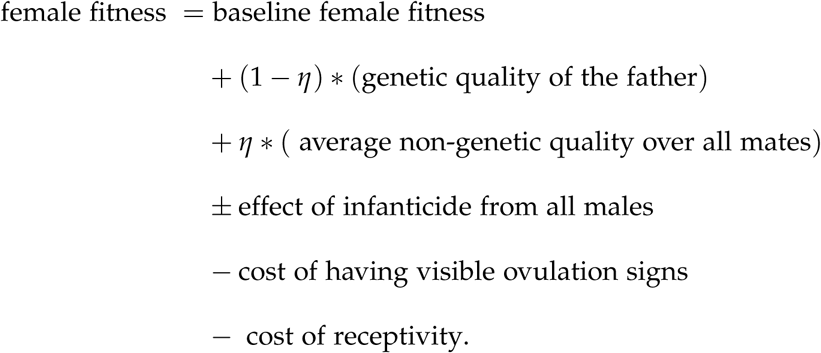

Our goal is to determine how female receptivity length evolves over time, both with and without visual ovulation signs simultaneously evolving. Appendix A provides further details, explicit fitness functions, and full definitions of equations and parameters.

## Results

We present our results in 3 different sets which differ in the trait(s) being selected:

1. Evolution of receptivity (*r*) given fixed concealed ovulation (*m*, *l* = 0)
2. Evolution of receptivity (*r*) given fixed ovulation signs present (*m* = 0.9, *l* = 4), and
3. Evolution of receptivity (*r*), ovulation signs magnitude (*m*), and ovulation signs length (*l*).

Note that studying the evolution of female sexual receptivity in isolation first (i.e., Sets 1 and 2) will allow us to better understand the coevolution of receptivity length with ovulation signaling. For each set, we investigate our model both with and without the effects of infanticide.

In all cases, numerical simulations of our model show that the average values of the receptivity trait (*r*) (and ovulation signs magnitude trait (*m*) and ovulation signs length trait (*l*), if included) converge to the unique equilibria *r^∗^* (and *m^∗^*, *l^∗^*, if included). A more complete investigation of the effects of all parameters can be found in the Online Appendix.

### Set 1: Evolution of receptivity (***r***) given fixed concealed ovulation (***m*, *l* = 0**)

Results from Set 1 can all be seen in the white bars of Fig. 2. The effects of *N*, *b*, and *c_r_* are visible in the white bars in Fig. 2(a). We intuitively expect that increasing the receptivity costs parameter (*c_r_*) will decrease receptivity length equilibrium *r^∗^*, and that is indeed the case. We also see that group size (*N*) and variation in male quality (*b*) have little effect on female receptivity length.

In the white bars of Fig. 2(b), we see increased receptivity length *r^∗^* with increased effects of infanticide (i.e., increased *α* and *β*). These results are intuitive since with stronger effects of infanticide come an increased benefit to females of mating with multiple males. Having a longer length of receptivity allows females more time to be able to mate with multiple males, which in turn makes multiple males think they may be the father of her offspring and hence help protect (or at least not harm) that offspring. Recall parameter *κ* controls how much males take visible ovulation signs into account when estimating their paternity. In the concealed ovulation case, decreasing *κ* results in increased receptivity length *r^∗^*. Even with concealed ovulation, if males only take the number of matings with a female into account when calculating their perceived probability of paternity, females are able to get benefit from mating at any point of their cycle. Hence there becomes an increased benefit to females for having longer lengths of receptivity.

Finally, Fig. 2(c) investigates the effects male traits have on female receptivity length. Increasing the relative weighting of male non-genetic effects (i.e., increasing *η*) increases *r^∗^*. As male non-genetic benefits become more valuable to females, females have increased incentive to mate outside of their fertility window, hence increasing their length of receptivity. *ρ*, the correlation between male genetic and non-genetic effects, has very little effect.

### Set 2: Evolution of receptivity (***r***) given fixed ovulation signs present (***m* = 0.9, *l* = 4**)

Results from Set 2 can all be seen in the black bars of Fig. 2. Looking at receptivity length under conditions of fully visible ovulation signs, we see the same effects as in Set 1 of parameters *c_r_*, *η*, *α*, and *β*, and the same non-effects of *N*, *b*, and *ρ*. The parameter *κ* also has the same effects as in Set 1 for similar reasons. As males put less weight in the amount of visible ovulation signs a female has when mating with her, females are able to get more benefit in mating with multiple males by mating at times of their cycle when they have fewer (or even no) signs visible. Hence there again becomes an increased benefit to females for having longer lengths of receptivity with smaller *κ*. These effects are all visible in the black bars of Fig. 2(a,c) (no infanticide) and Fig. 2(b) (with infanticide).

**Figure 2:**
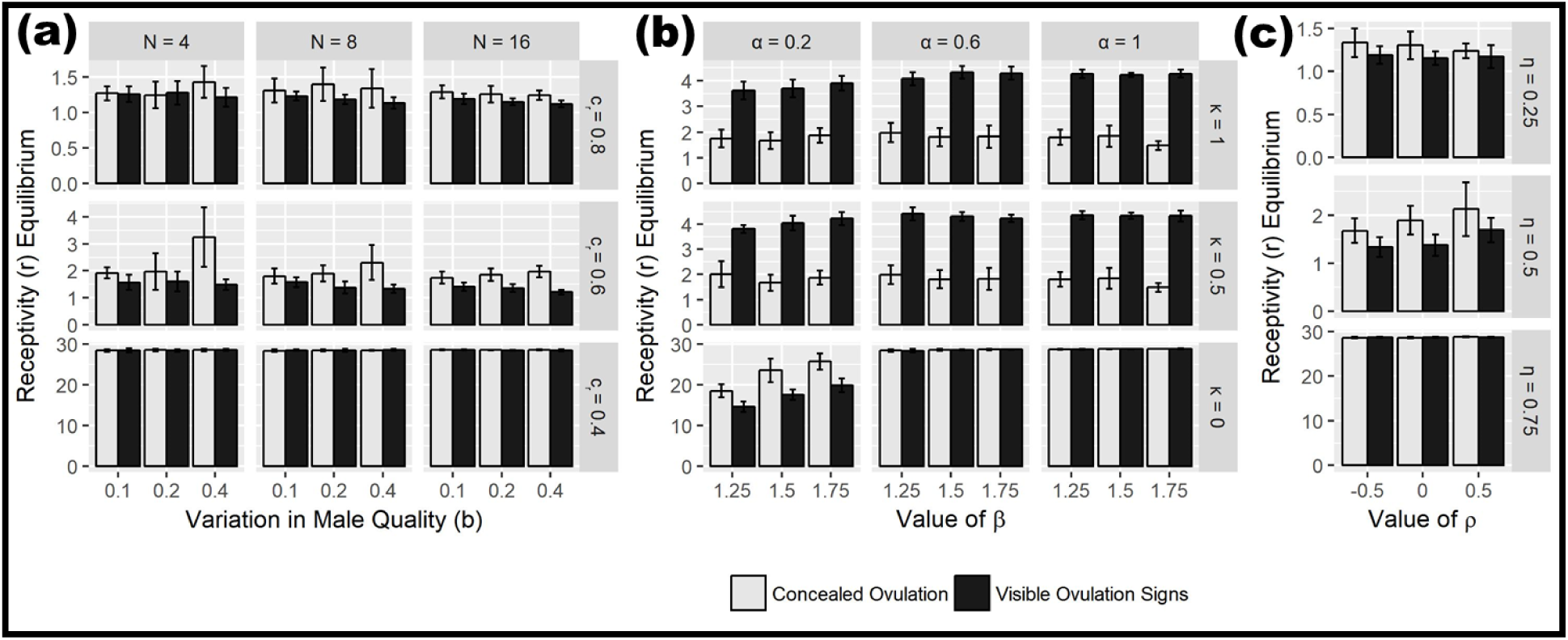
The effects of varying parameters on the average equilibria values of receptivity (*r^∗^*) in the case when only *r* is allowed to evolve. Note the different y-axis scales. White bars indicate Set 1 (assuming concealed ovulation: *m*, *l* = 0), while black bars indicate Case 2 (assuming fixed visible ovulation signs: *m* = 0.9, *l* = 4). Equilibria are obtained by averaging over 10 initial condition runs (with standard deviation indicated by error bars). (a) Varying *N* (group size), *b* (variation in male quality), and *c_r_* (costs of receptivity) in the case of no infanticide (*α* = 0, *ρ* = 0, *η* = 0.5). (b) Varying *α* (maximum benefit of infanticide protection), *β* (maximum cost of infanticide), and *κ* (weight males put on visible ovulation signs in estimating paternity) in the case of with infanticide (*N* = 8, *b* = 0.2, *c_r_* = 0.6, *ρ* = 0, *η* = 0.5). (c) Varying *η* (relative weighting of male non-genetic effects) and *ρ* (correlation between male genetic and non-genetic effects), in the case of no infanticide (*N* = 8, *b* = 0.2, *c_r_* = 0.6, *α* = 0). Other parameters held constant throughout: *c* = 0.2, *∊_m_* = 0.25, *∊_f_* = 0.01.

Note that extended female sexual receptivity length can evolve both with *and* without visual ovulation signs being present (i.e., in both Set 1 and Set 2). This is interesting in that neither concealed ovulation nor visual ovulation signs are necessary for female receptivity lengths to evolve. Indeed, when continuous receptivity is most beneficial (for example, with low costs of receptivity *c_r_*, strong effects of infanticide *α*, *β*, and/or increased weight placed on non-genetic male benefits *η*), continuous receptivity evolves in both the concealed ovulation and visible ovulation signs cases. When continuous receptivity is less beneficial, we see receptivity length *r^∗^* typically not exceed 4 when visible ovulation signs are present. This is due to ovulation signs length *l* being fixed at 4, meaning receptivity length is typically not evolving to be longer than the amount of time the female has ovulation signs visible, an intuitive result.

### Set 3: Evolution of receptivity (***r***), ovulation magnitude (***m***), and ovulation length (***l***)

We now allow for each of receptivity length (*r*), magnitude of visible ovulation signs (*m*), and length of visible ovulation signs (*l*) to independently evolve. We compare these results with not just each other, but also with results for *m^∗^* and *l^∗^* when *r* does not evolve and continuous receptivity is instead assumed. Note full results for the evolution of *m* and *l* with continuous receptivity assumed can be found in Rooker and Gavrilets (2018). Our results for Set 3 are shown in Fig. 3 (a, c, without infanticide and b, with infanticide). In Fig. 3(a) we see the costs of having visual ovulation signs *c* to have little effect on receptivity length *r^∗^*and to have similar effects as in Rooker and Gavrilets (2018) on *m^∗^* and *l^∗^*, both as expected. In Fig. 3(b) we see receptivity length equilibria (*r^∗^*), ovulation signs magnitude equilibria (*m^∗^*), and ovulation signs length equilibria (*l^∗^*) increase with the impacts of infanticide *α* and *β*. As discussed above and in Rooker and Gavrilets (2018), these results are intuitive.

**Figure 3:**
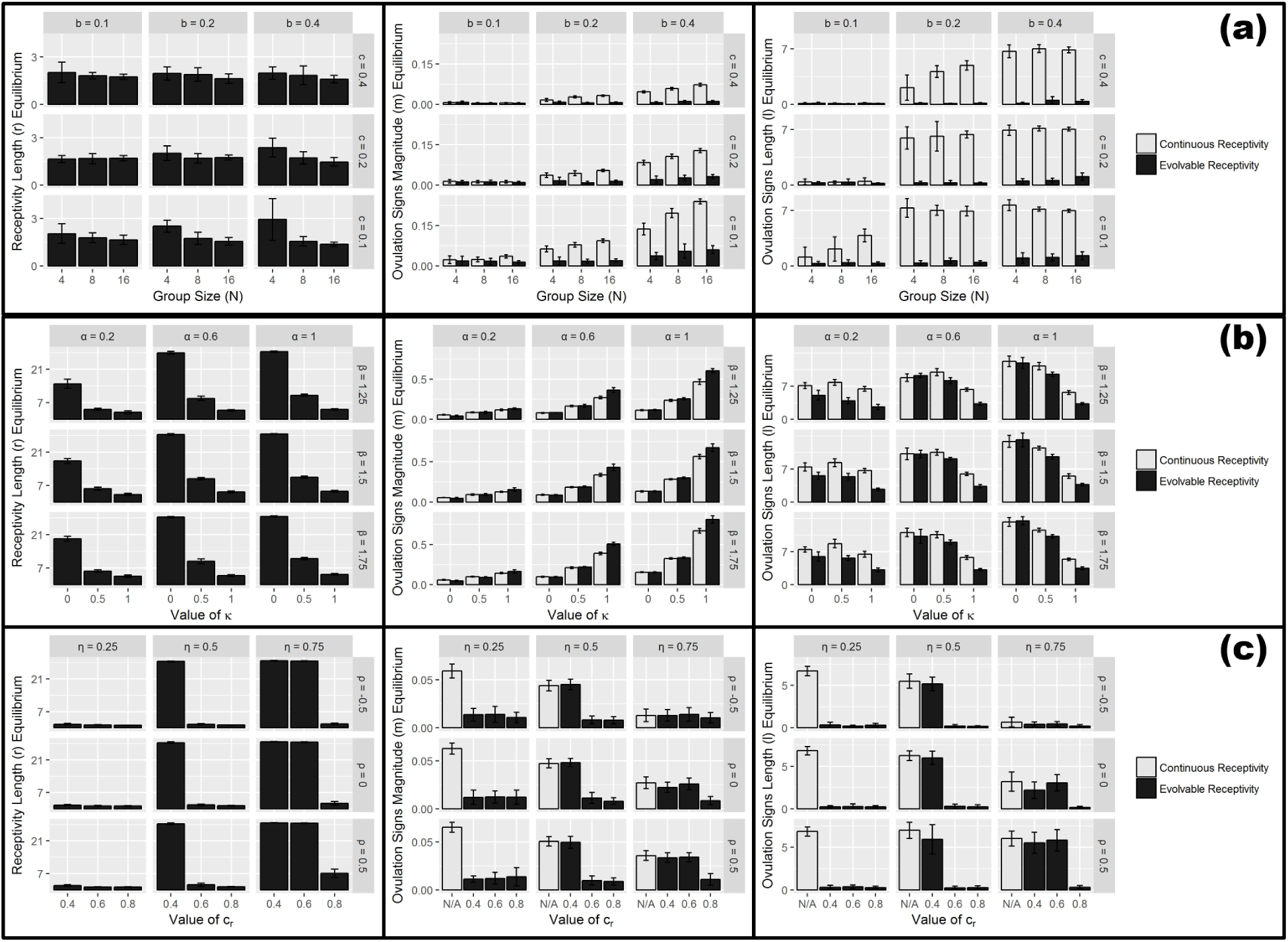
The effects of varying parameters on the average equilibria values of receptivity length (*r^∗^*, left graphs), ovulation signs magnitude (*m^∗^*, middle graphs), and ovulation signs length (*l^∗^*, right graphs), with both continuous receptivity assumed (white bars) and evolvable receptivity (black bars). Equilibria are obtained by averaging over 16 initial condition runs (with standard deviation indicated by error bars). (a) Varying *N* (group size), *b* (variation in male quality), and *c* (costs of having visible ovulation signs) in the case of no infanticide (*c_r_* = 0.6, *α* = 0, *ρ* = 0, *η* = 0.5). (b) Varying *α* (maximum benefit of infanticide protection), *β* (maximum cost of infanticide), and *κ* (weight males put on visible ovulation signs in estimating paternity) in the case of with infanticide (*N* = 8, *b* = 0.2, *c_r_* = 0.6, *ρ* = 0, *η* = 0.5). (c) Varying *η* (relative weighting of non-genetic effects), *ρ* (correlation between male genetic and non-genetic effects), and *c_r_* (costs of receptivity) in the case of no infanticide (*N* = 8, *b* = 0.2, *c* = 0.2, *α* = 0.6). All other parameters held constant: *∊_m_* = 0.25, *∊_f_* = 0.01.

We also begin to see a trade-off between receptivity length and ovulation signs magnitude. For example, in Fig. 3(b), we see that increasing *κ* (parameter controlling how much males take visible ovulation signs into account when estimating their paternity) increases *m^∗^* and decreases *r^∗^*. This pattern is due to the importance of paternity concentration. The stronger the magnitude of the ovulation signal, the more beneficial it is to the female to only be receptive around those day(s) of having a peak signal, thus concentrating paternity in high-quality males.

In the extreme case, when *κ* = 0, males are only taking the *number* of matings with a female into account when estimating their paternity, thus ignoring any visible ovulation signs present for their paternity estimate (but still using such signs to find the most likely fertile mates). As the effects of infanticide increase, females gain increasing benefit from mating with multiple males during any one cycle. With *κ* = 0, a female is able to make males think they might be the father of her offspring by mating with them even when she has zero ovulation signs visible. Hence it becomes advantageous for obtaining multiple matings (and female fitness, provided the effects of infanticide are large enough) to have as long a length of receptivity as possible, in order to be able to mate with the most males possible.

In Fig. 3(c), we see that increasing *ρ* (the correlation between male genetic and non-genetic effects) slightly increases each of receptivity length and ovulation signs. Increasing *ρ* increases the value to females from mating with high-quality males since such high-quality males now effectively have ‘double’ the quality (i.e., genetic and non-genetic). In line with the trade-offs discussed earlier, we also see that increased *η* (the relative weighting put on non-genetic vs. genetic effects in males) results in increased receptivity length (since females have more incentive to mate outside their fertility window with higher non-genetic effects) and decreased visible ovulation signs (since attracting high-genetic quality males becomes less important).

We also compared these results to the case when receptivity is assumed to be continuous (i.e., females are willing and able to mate on every day of their cycle). The continuous receptivity results are displayed as the white bars in Fig. 3, and the evolvable receptivity results as the black bars in Fig. 3. In general, introducing evolvable receptivity strongly decreases *m^∗^* and *l^∗^* in the case of no infanticide, and increases *m^∗^* and slightly decreases *l^∗^* in the case of with infanticide. With evolvable receptivity, females are able to concentrate paternity by having a short length both of receptivity *and* of visible ovulation signs, a pattern we see when infanticide is not present. Then considering the effects of infanticide, paternity confusion becomes increasingly important to females in addition to paternity concentration, leading to those longer lengths of visible ovulation signs. In addition, since infanticide increases the stakes of the female-female competition, females are now able to devote more resources towards strong ovulation signs magnitude for those fewer days of their receptivity, the precise pattern we see in Fig. 3(b).

Given the large number of parameters, visual inspection of graphs is not enough. Therefore, for both with and without infanticide in Set 3, we also run an analysis of variance (ANOVA) to determine which of receptivity length *r*, ovulation magnitude *m*, and ovulation length *l* are affected most by which parameters. Full results of these tests can be found in the Online Appendix. In general, what we find in the without infanticide case, is that (intuitively) costs of receptivity *c_r_* mainly affects *r*. Conversely, costs of having visible ovulation signs *c* only affects *m* and *l*. We also find that the benefit of male quality *b* strongly increases both *m* and *l*, while having no effect on *r*. Group size *N* increases *m*, but has no effect on *r* or *l*. Increasing *η* (the relative weighting of male non-genetic effects) increases each of *r*, *m*, *l*, but has the largest effect on *r*. Finally, *ρ* has very little effect on any of *r*, *m*, *l*.

In general for the with infanticide case, we find that *β* has little effect. When *β* does have an effect, such an effect is always positive. On the other hand, *α* positively affects each of *r*, *m*, *l*, although *α* affects *m*, *l* more so than *r*. *κ* strongly increases *m* and strongly decreases both *l* and *r*. Note that all of these results reaffirm the patterns discussed above.

With regards to the reproductive stochasticity parameters *∊_m_* and *∊_f_*, we find no effects on receptivity length *r*. As in Rooker and Gavrilets (2018), increasing *∊_m_* will result in decreased *m^∗^*, *l^∗^* (since increasing *∊_m_* makes the ‘prize’ for females of winning the female-female competition less valuable).

## Discussion

It was our goal here to investigate theoretically the evolutionary forces regarding female primate sexual receptivity. In particular, we were interested in under what conditions were continuous receptivity and/or very short lengths of receptivity able to evolve. We were also interested in the effects infanticide could have on the evolution of receptivity, as well as how the evolution of sexual attractiveness (visual ovulation signs vs. concealed ovulation) could affect the evolution of receptivity. Our model has produced four main results of interest in answering our research questions.

First, how does length of receptivity contribute to female reproductive success? Female reproductive success here can be thought of as a trade-off between paternity concentration and paternity confusion (van Schaik et al., 1999). A female concentrating paternity in high-quality males allows her offspring to have higher quality, and hence increases the female’s fitness. Conversely, a female mating with multiple males during one cycle will confuse paternity, i.e., make multiple males think they may be the father of her subsequent offspring. Paternity confusion can increase a female’s fitness, for example by reducing her offspring’s risk of infanticide. Mating with multiple males could also increase a female’s fitness via other non-genetic benefits, such as protection or provisionings.

Our modeling work shows this trade-off is manifested in females’ length of receptivity. In situations where paternity concentration is favored, short lengths of receptivity are seen. Conversely, in situations where paternity confusion is favored (e.g., when effects of infanticide are strong), longer lengths of receptivity are seen. Short lengths of receptivity allow females to only mate on days where they have strong visual ovulation signals, meaning on those days where they are able to attract the highest-quality mates. By only mating on those few days of peak signal, such females are able to concentrate paternity of their offspring in high-quality males. On the other hand, long lengths of receptivity give females ample time to mate with as many males as possible, thus allowing them to confuse paternity or gain increased non-genetic benefits. It is in this way that receptivity is able to contribute to both paternity concentration and paternity confusion, and hence also female reproductive success.

Second, what role could infanticide have in the evolution of receptivity? When infanticide is introduced, our model shows that receptivity increases as the effects of infanticide get stronger (i.e., larger *α*, *β*). Introducing the threat of infanticide increases the importance of paternity confusion, and as explained above, longer lengths of receptivity allow for more matings with multiple males, meaning increased paternity confusion. Thus, increasing the effects of infanticide increases the equilibria length of receptivity.

Under conditions of infanticide, we also find that increasing the importance for males estimating their paternity of the female having signs present when mating leads to decreased receptivity length. Note this is equivalent to increasing the parameter *κ*, i.e., moving away from males estimating their paternity from simply mating with a female. Increasing *κ* increases the weight males put on a female’s ovulation signs when evaluating their estimated paternity. This means with increased *κ*, males are putting *more* weight on females having visual ovulation signs present, which means simply mating with more males may not benefit the female at all; it may only be of benefit to mate when she *does* have strong ovulation signs visible, leading to increasing *κ* decreasing equilibria receptivity length.

Third, what factors facilitate the evolution of continuous receptivity? Our model shows three situations which can easily lead to continuous receptivity: (1) decreased costs of receptivity (*c_r_*), (2) increased weighting of non-genetic male effects (*η*), and (3) increased effects of infanticide (*α*, *β*) and in particular *decreased* weighting of ovulation signs when males estimate their paternity (*κ*). Note the first two can happen even without infanticide present at all.

Without infanticide, the only benefit to females from mating is coming from the quality (genetic or otherwise) of the males with whom she mates. In particular, continuous receptivity is more likely to evolve under these conditions with lower costs of receptivity (*c_r_*) and increased weighting of male non-genetic effects (*η*). The influence of *c_r_* is quite intuitive (i.e., lower costs make its evolution more likely), as is *η* (i.e., increased weighting on male non-genetic benefits means females can mate outside of their fertile window and still receive such benefits).

Continuous receptivity can also evolve under conditions of infanticide, regardless of whether ovulation signs are visible or not. As explained earlier, *κ* is an important parameter influencing receptivity length. At its extreme, *κ* = 0, males are not putting *any* weight on a female’s visual ovulation signs; rather, the male when estimating his paternity of an offspring takes into account *only* the number of matings with its mother during that cycle. In this case, with *κ* = 0, a female is able to confuse paternity throughout her entire length of receptivity, regardless of when, if, or how many visual ovulation signs she has present. Hence, when *κ* = 0, as expected we can see continuous receptivity evolve.

Fourth, how is receptivity linked to the evolution of sexual attractiveness? Recall that in the case of fixed visible ovulation signs (i.e., with only *r* evolving, see Fig.2), we postulated visible ovulation signs to have a fixed length of 4. Other than the continuous receptivity cases discussed above, we do indeed see receptivity length equilibria less than or equal to 4. When receptivity is instead allowed to evolve, receptivity is evolving to mirror the ovulation signs length, when infanticide is present. Conversely, without infanticide present, there is no benefit to females of paternity confusion, meaning it is now in females’ best interests to concentrate paternity by only mating for a very short portion of their cycle (i.e., those 1-2 days).

In general, with small *η*, we do not see receptivity length longer than a female’s fertility length. This makes sense since outside of the continuous receptivity cases discussed earlier, with a large enough cost (*c_r_*) associated with being receptive, females should only be receptive during the portion of their cycle where they are actually able to conceive. In turn, we also typically see a female’s ovulation signs length equilibria match her receptivity length equilibria, meaning both receptivity and a female’s length of ovulation signs being visible will typically not be longer than her length of fertility. Such continuity between receptivity length and ovulation signs length is intuitive, since again, females are generally expected to only spend energy on advertising their fertility when they are indeed receptive to mating. However, with a large enough *η*, it becomes in females’ best interest to instead have very long lengths of receptivity, in order to extend the amount of time in which they can receive all the non-genetic benefits from mating with males (even if not conceiving).

Finally, there also appears to be a trade-off between receptivity length and ovulation signs magnitude (see Fig. 3). When each of receptivity (*r*), ovulation signs magnitude (*m*), and ovulation signs length (*l*) are allowed to evolve, we find that with long lengths of receptivity (i.e., at or near being continuously receptive), ovulation signs magnitude (*m*) is very small. This is suggestive of the possible relationship between concealed ovulation and continuous receptivity, as is the case in humans. Such a result is most likely to occur with large *η*, small *c_r_*, small *ρ*, and no infanticide. This means the most favorable conditions for concealed ovulation and continuous receptivity to evolve together are: (1) Large non-genetic benefits from males, (2) Small costs to females of being sexually receptive, (3) Negative correlation between genetic and non-genetic benefits from males, and (4) No infanticide present.

In contrast, we also see cases where short lengths of receptivity evolve with very strong visible ovulation signs. This is most likely to occur when infanticide is present and its effects are very strong (i.e., large *α*, *β*). In addition, *κ* must also be large, meaning males are estimating their paternity using the amount of ovulation signs visible during their matings with any females.

Our approach comes with several limitations. Like many models, we assume discrete generations for mathematical simplicity. We also assume an additive fitness function. Explicitly accounting for the female cycle has made obtaining informative analytical results not feasible, leading to us relying on agent-based simulations. In our model, we did not impose female cycle synchrony. Assuming so would increase female-female competition and hence ovulation signaling and receptivity. We do not consider the effects of any extended mate-guarding (i.e. mate-guarding lasting longer than our one discrete unit of time). We do not explicitly allow for variation in signal reliability, instead assuming a female’s peak day of fertility and day of peak visible ovulation signs to line up. We also only consider multi-male, multi-female mating systems, excluding other mating systems from direct consideration in our model. In addition, our description of ‘male quality’ is purposefully general; we do not explicitly take into account male age, rank, or any other similar known factor in primate mating.

While much work still needs to go into understanding the evolution of sexual receptivity in female primates, our modeling suggests several pathways by which longer receptivity could have evolved or diminished. While it was our hope to test our model’s predictions on the limited data available on female primate sexual receptivity length (e.g., see Hrdy and Whitten (1987); Stockley (2002); van Schaik et al. (1999)), such data turned out to be too sparse, especially when only considering polygynandrous mating systems as does our model, and also at times variable between sources. In addition, there can be intraspecific variation in receptivity length as well, which our model does not specifically address (for example, receptivity lengthening when a new male takes over a group) (van Schaik et al., 1999). More empirical data and further detailed information on fitness components and ecological factors would be needed in order to do an effective comparative empirical analysis, which we hope our work to have inspired.

## Appendix A: Methods Details

### Model Set-up

We consider a population of individuals interacting in *G* groups, each comprised of *N* males and *N* females. Male-provided genetic (*y_g_*) and non-genetic (*y_ng_*) benefits are randomly drawn from the bivariate normal distribution, each with mean 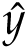 and standard deviation *b*, and correlation parameter *ρ*; parameter 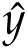 characterizes mean male quality and *b >* 0 the extent of variation. Males are ranked according to the value of their genetic quality: *y_g,j_* such that *y_g_*_,1_ *> y_g_*_,2_ *>* … for each male *j*. Small *b* implies small additional benefits to females from investing in ovulation signaling. For simplicity, the distribution of male quality in the population remains constant, e.g., as a result of mutation-selection balance.

Our model explicitly accounts for the female cycle by using *D* discrete units of time. Without loss of generality, we refer to these units of time as ‘days’ (e.g., *D* = 29 days). We assume each female to mate once on every day on which she is receptive, and each male to be able to mate with no more than one female on any given day. Biologically, a male after mating with a female may guard her to ensure no other male is able to mate with her (i.e., mate guard) and/or a male must do other activities besides mating/searching for mates (e.g., hunting/eating). Mating is followed by offspring production and dispersal.

For each day of the cycle, every female will have an associated probability of fertilization. We assume each fertile period to last *C ≤ D* days (e.g., *C* = 7). For all days lying outside these *C* fertile days, females are assumed to have zero probability of fertilization but can still mate if receptive on those days. We assume a triangular shape for the fertility function to be identical in all females, but with midpoints randomly distributed so as cycle synchrony occurs only probabilistically. Visual ovulation signs and receptivity are both correlated with these days of fertility (in that a female’s day of peak ovulation signs will align with both her median day of receptivity and her peak day of fertility), but can each still evolve freely.

Let *r* be the number of days of the cycle a female is receptive to mating (a non-negative, integer value). We treat visual ovulation signs, *x*(*d*) for each day *d* of the cycle, as overlapping graded curves. Each female’s curve is characterized by two traits: magnitude (*m*) and length (*l*). *m* is the maximum amount of ovulation signs a female has visible during her cycle (a nonnegative, continuous value), while *l* is the number of days a female has *some* amount of ovulation signs visible (a non-negative, integer value).

Given female traits *m_i_ ≥* 0 (ovulation signs magnitude) and *l_i_ ∈* Z, *l_i_ ∈* [0, *D*] (ovulation signs length), the amount *x_i_*(*d*) of ovulation signs visible on any day *d* of the cycle is defined as

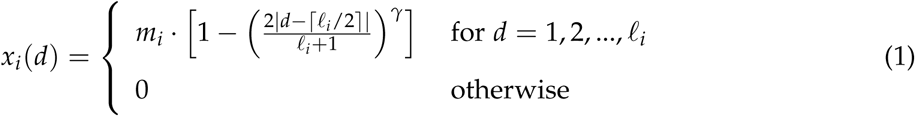

for parameter *γ >* 0 and *d* measured from the start of ovulation signs being visible. Note *m_i_* represents the peak of *x_i_* (*d*) and *f_i_* the width of the non-zero portion of *x_i_* (*d*). Mutation effects in *m_i_* are randomly chosen from the normal distribution *N*(0, *σ*^2^), whereas mutations in *f_i_* and *r_i_* are discrete, representing adding or subtracting exactly one day to or from a female’s time of having ovulation signs visible or being receptive, respectively.

On each unit of time, mating pairs are formed by first randomly perturbing both the male trait *y* and female trait *x* by adding to each an independent, normally distributed random variable with standard deviation *∊_m_* and *∊_f_*, respectively. Males and receptive females are then sorted according to these perturbed values and mating occurs between individuals of the same order. This means if there are *R* receptive females on any given day of the cycle, there will only be *R* males able to mate with a female on that day.

Selection occurs for males via mating success and for females via fertility selection. In the case of no infanticide, we define female fitness (fertility) as

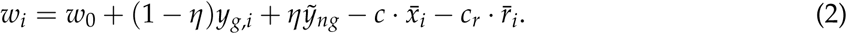

Here, *w*_0_ is baseline fitness, *y_g,i_* the genetic benefit provided by the male who fertilizes female *i*, the 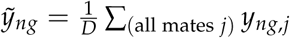 average non-genetic benefit provided by all female *i*’s mates, *c · x̄_i_* the costs to a female of supporting her visual ovulation signs with 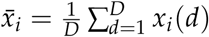, a female’s average visual ovulation signs, and *c_r_· r̄_i_* the costs to a female of receptivity with 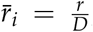, a female’s proportion of the cycle being receptive.

Each male in a female’s group can affect her fitness directly, due to infanticide, although males will differ in their actions’ effectiveness (e.g., due to strength, rank, size, alliances, etc.). We assume each male’s effectiveness to be proportional to *f* (*j*) for each male *j* (sorted by males’ *y_g_*) via an exponential function: *f* (*j*) *∼ e^−ωj^* with parameter *ω >* 0 controlling the amount of disparity among males in their corresponding effectiveness within the group. Note a larger value of *ω* indicates more disparity, and *ω* = 0 equality.

A male’s contribution *g*(*p_j_*) to female fitness (positive via helping protect the offspring, or negative via not helping and/or harming the offspring) depends on his perceived probability of paternity *p_j_*. Following Rooker and Gavrilets (2018), we define 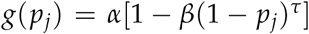, with parameters *α*, *β >* 0 and *τ ≥* 1. Note *α* determines the maximum benefit a female can obtain from a male protecting her offspring from infanticide, while *α*(1 *− β*) determines the maximum cost a female can incur from a male *not* protecting her offspring from infanticide. The overall effect of infanticide on female fitness (that gets added to the right-hand side of Eq. 2) is ∑*_j_ f* (*j*)*g*(*p_j_*), summing over all males *j*. Note whenever *α* = 0, there are no effects of infanticide.

A male in general would not know his actual paternity probability and would instead have to estimate it, when decided on protecting an infant or committing infanticide. Consider male *j* mating with female *i* with visible ovulation signs *x_i_*(*d*). We postulate that the male-estimated (“perceived”) probability *p_j_* of being the father of her offspring is proportional to (*x_i_*(*d*))*^κ^* where *κ ∈* [0, 1] is the weight males put on visual ovulation signs. With multiple matings with the same female, we sum up the corresponding terms (*x_i_*(*d*))*^κ^*. More specific model details can be found in the Online Appendix.

### Simulations

This model was implemented in C++. Migration occurs with only females migrating between groups, although simulations were also run with only males migrating between groups and the results were unaffected. Mutation occurs with a mutation rate per gene per generation of 10*^−^*^3^. Mutations for *m* are selected from N(0, 0.1), while mutations for *l* and *r* independently from the set *{−*1, 1*}* with equal probability. 2*N* offspring are created in each group every generation, with *N* female and *N* male, determined randomly. The expected number of offspring per female is 2; the actual number is random, proportional to each female’s fitness.

For this model, we consider 3 different sets regarding selection:

1. Evolution of receptivity (*r*) given fixed concealed ovulation (*m*, *l* = 0)
2. Evolution of receptivity (*r*) given fixed ovulation signs present (*m* = 0.9, *l* = 4), and
3. Evolution of receptivity (*r*), ovulation magnitude (*m*), and ovulation length (*l*).

For (1) and (2), we use the following 10 initial conditions: *r* = 1, 4, 7, 10, 13, 16, 19, 22, 25, 28. For (3)’s initial conditions, we use every combination of *r* = 3, 11, 19, 27, *m* = 0.3, 0.9, and *l* = 2, 4. Parameters used for all the above simulations include: *G* = 400; *T* = 100, 000; *D* = 29; *C* = 7 (with probabilities of fertilization on each of these days being 0.125, 0.25, 0.375, 0.5, 0.375, 0.25, 0.125, respectively); *w*_0_ = 2; *N* = 4, 8, 16; *b* = 0.1, 0.2, 0.4; *c* = 0.1, 0.2, 0.4; *c_r_* = 0.4, 0.6, 0.8; *η* = 0.25, 0.5, 0.75; *ρ* = *−*0.5, 0, 0.5; *γ* = 1; 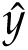 = 1; 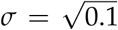; *∊_m_* = 0, 0.25, 0.5; *∊_f_* = 0, 0.01, 0.05, *α* = 0, 0.2, 0.6, 1; *β* = 1.25, 1.5, 1.75; *τ* = 2; *ω* = 1; and *κ* = 0, 0.5, 1. Further information on parameter values tested and each of their effects is available in the Online Appendix.

## Acknowledgments

We thank Susan Perry and Sarah Hrdy for comments and suggestions. K.R. was supported by a National Science Foundation Graduate Research Fellowship. S.G. was supported by the National Institute for Mathematical and Biological Synthesis through National Science Foundation awards EF-0832858 and DBI-1300426, with additional support from The University of Tennessee.

## Online Appendix

### Supplementary Information for

#### Further Details of the Model

##### The Female Cycle

Female estrous cycles are divided into *D* discrete units of time (e.g., *D* = 29 days) with *C ≤ D* days of fertility (e.g., *C* = 7). Although the form of such fertile periods will be identical in all females, the timing of each fertile period will be randomly distributed among females. For instance, when *C* = 7 (calling each of these fertile days *C*1, *C*2, …, *C*7) and *D* = 29 (calling each of these days *D*1, *D*2, …, *D*29), *C*1 in every female has an equal probability of landing on any of *D*1, *D*2, …, *D*29. Since *C* is a cycle, we similarly assume *C*7 can land on any such day. For example, if *C*1 landed on *D*24, then *C*7 would ‘wrap around’ the cycle to land on *D*1. Moreover, for all females we assume ovulation happens directly in the middle of this *C* cycle. Hence, for *C* = 7 we assume ovulation happens on *C*4 and the probabilities of fertilization increase from *C*1 to *C*4 and then decrease from *C*4 to *C*7.

Each female *i* will have *r_i_ ≤ D* days of receptivity. These days of receptivity line up with the female’s days of fertility, such that a female’s median day of fertility will equal her median day of receptivity. Similarly, since having visible ovulation signs at least loosely correlates with a female’s fertility, we assume the day(s) of having maximum visual ovulation signs also align with the day(s) of that female’s maximum fertilization probability, and align these cycles accordingly (note in most cases, *C ≠ r_i_ ≠ l_i_*).

Recall that *x_i_* (*d*) denotes the amount of visual ovulation signs present on each day *d* of the female cycle, with *x_i_*(*d*) defined by the function

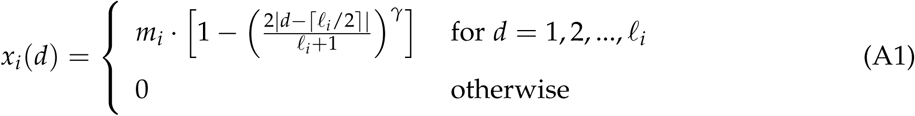

This function is determined by *m_i_* (ovulation signs magnitude), *l_i_* (ovulation signs length), and parameter *γ* (controlling the shape of the resulting curve). Note the exact function *x_i_*(*d*) is chosen such that (i) *m_i_* will denote the peak of the curve, (ii) *lremain independent of each other,_i_* will denote the width of the non-zero portion of the curve, (iii) *m_i_*, *l_i_* remain independent of each other, and (iv) *m_i_*, *l_i_* can both decrease all the way to zero. Example values of *r_i_* are displayed in Fig. 1 via the light blue shading, for *m_i_* = 1, *f_i_* = 7, and *C* = 7. Ovulation signs on each day are depicted by the red shaded bars, while fertility probabilities are given by the black lines.

##### Determining Male-Female Mate Pairs

We assume that, for every day of the cycle, males of higher GC will preferentially mate with females with more ovulation signs visible. We also assume each individual to mate at most once on each day of the cycle. Females will only mate on days on which they are receptive, and males will only mate if there is a receptive female with whom to mate. Parameters *∊_m_* and *∊_f_* control the amount of stochasticity in the process of mating pair formation. On each day, mating pairs are formed by first randomly perturbing both the male trait *y_g_* and female trait *x* by adding to each an independent, normally distributed random variable with standard deviation *∊_m_ ≥* 0 and *∊_f_ ≥* 0, respectively (i.e., *′* = *x* + *∊_x_*, *y′_g_* = *y_g_* + *e_y_*, where 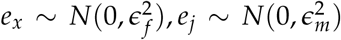). Males and receptive females are then sorted according to these perturbed values and mating occurs between individuals of the same order. With *∊_m_*, *∊_f_ →* ∞, mating pairs are formed completely independently of the values of *x_i_* and *y_g,j_*.

**Figure 1:**
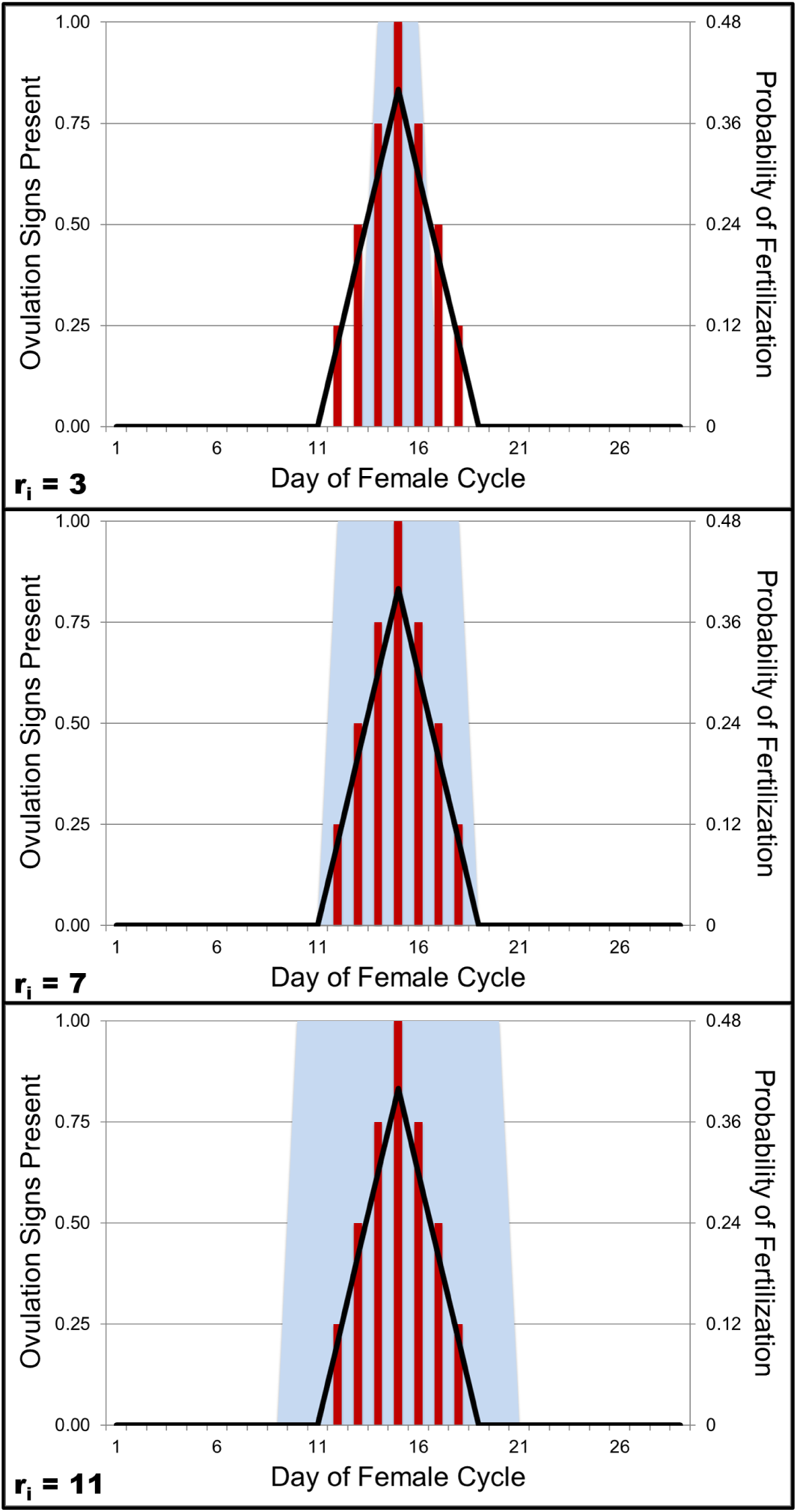
Sample receptivity lengths are displayed for *r_i_* = 3, 7, 11 (from top to bottom) via the light blue shading. Each graph also includes ovulation signs present (with *m_i_* = 1, *l_i_* = 7) via the red bars and fertilization probabilities (with *C* = 7) via the black lines. In the top graph, we see that a female can be fertile and/or have ovulation signs visible on days where she is not receptive. In the bottom graph, we instead see a female can be receptive on days where she does not have any ovulation signs visible and/or is not fertile. The middle graph depicts the situation where visible ovulation signs, fertility, and receptivity all line up with lengths of 7 days.

##### Calculating Probabilities of Paternity

To calculate the probability of paternity for each male, first the fertility of each female with which that male has mated is summed up for every day a particular male mates with her. These values are normalized across males for each female in the group, meaning a male who mates with the female on a day where she has a higher fertility probability will have a higher paternity probability. The actual father for that female’s offspring is then determined randomly, proportional to each male’s probability of paternity for her offspring.

Note these *actual* probabilities of paternity are different from the *perceived* probabilities of paternity, as detailed in the main text. We make this distinction because males will not know any female’s probability of fertility at the time he mates with her; a male will only know how many visible ovulation signs she has present. Separating these quantities allows *actual* probabilities of paternity (which males do not know) to be calculated using females’ probabilities of fertility, and *perceived* probabilities of paternity (which males *do* know) to be calculated using females’ visible ovulation signs.

##### Infanticide

We define the exponential function of male effectiveness to be:

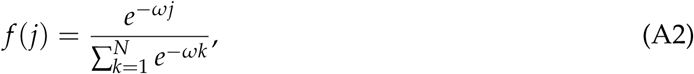

with parameter *ω >* 0 controlling the amount of disparity among males in their corresponding effectiveness within a group. Recall a larger value of *ω* indicates more disparity among males.

We define the function for male contributions to offspring survival to be

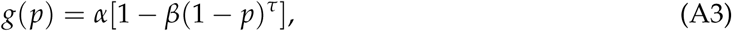

with parameters *α*, *β >* 0 and *τ ≥* 1. *α* determines the maximum benefit a female can obtain from a male protecting her offspring from infanticide, while *α*(1 *− β*) determines the maximum cost a female can incur from a male *not* protecting her offspring from infanticide. Note whenever *α* = 0, there are no effects of infanticide on infant survival.

##### Female Fitness Functions

The fitness function for female *i* in the model is as follows:

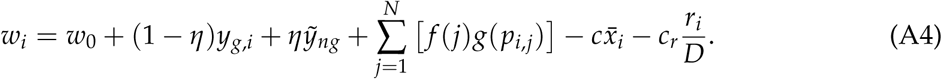

Here, *w*_0_ is baseline fitness, *y_g,i_* the genetic benefit provided by the male who fertilizes female *i*, 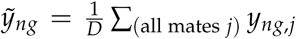 the average non-genetic benefit provided by all female *i*’s mates, *p_i,j_* the perceived paternity probability of male *j* for female *i*’s offspring, *c · x̄_i_*the costs to a female of supporting her visual ovulation signs with 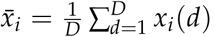, a female’s average visual ovulation signs, and 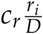 the costs to a female of being receptive to mating for 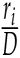 proportion of her cycle.

Notice for the case when *α* = 0 and, thus, *g* = 0 (i.e., without the effects of infanticide), the fitness function above collapses into

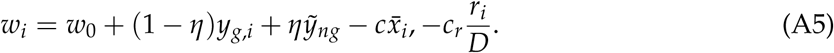

Note in this *α* = 0 (no infanticide) case, parameters *β*, *κ*, *τ*, and *ω* all become unnecessary.

#### List of all Parameters

Parameters used in the model are outlined in Table A1. For each parameter, one value listed implies the parameter remained fixed throughout all simulations, and multiple values indicates each of those parameter values was tested in all combinations with other parameters.

**Table A1:**
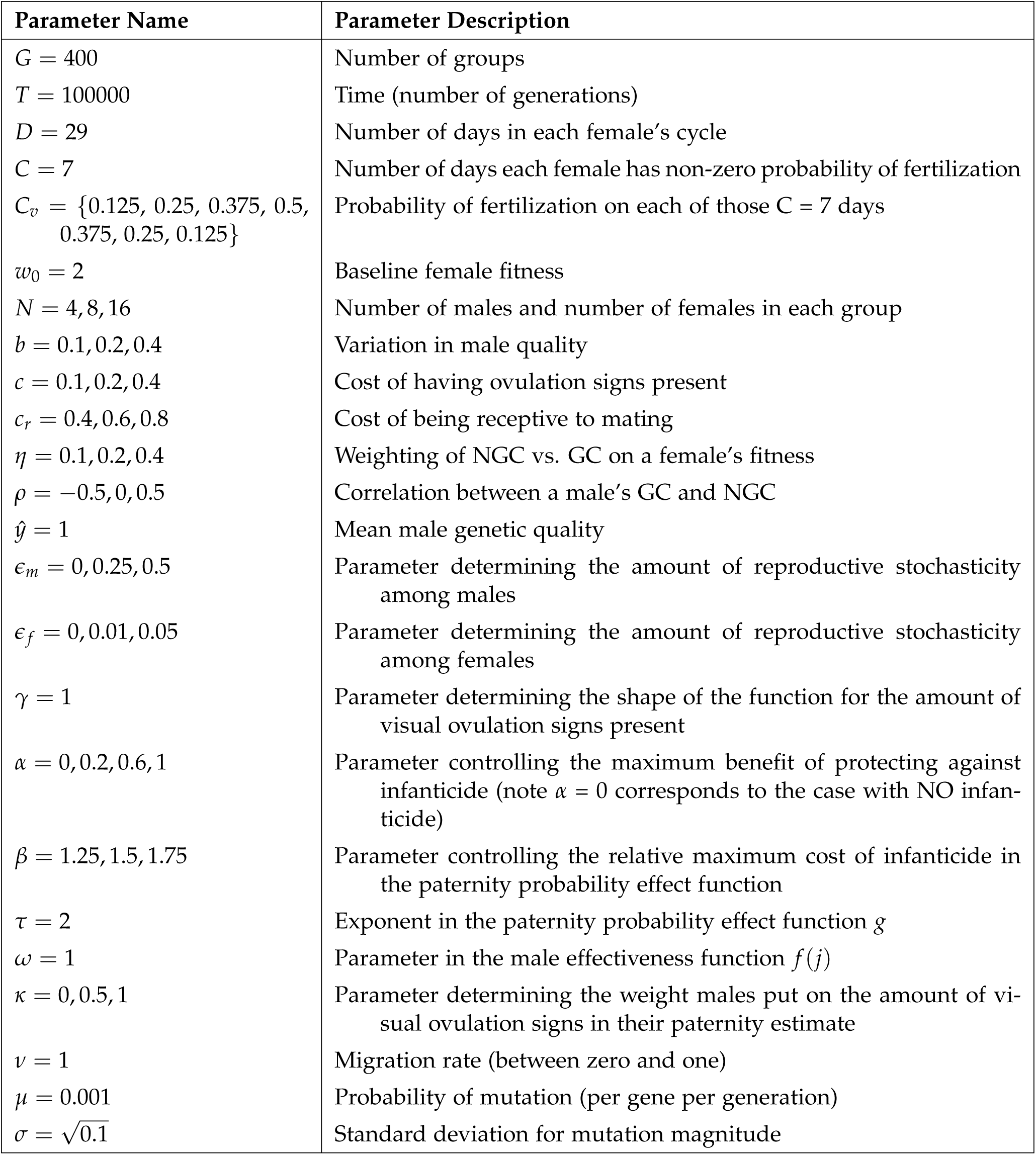
List of All Parameters

| Parameter Name | Parameter Description |
| --- | --- |
| $G = 400$ | Number of groups |
| $T = 100000$ | Time (number of generations) |
| $D = 29$ | Number of days in each female's cycle |
| $C = 7$ | Number of days each female has non-zero probability of fertilization |
| $C_v = \{0.125, 0.25, 0.375, 0.5, 0.375, 0.25, 0.125\}$ | Probability of fertilization on each of those $C = 7$ days |
| $w_0 = 2$ | Baseline female fitness |
| $N = 4, 8, 16$ | Number of males and number of females in each group |
| $b = 0.1, 0.2, 0.4$ | Variation in male quality |
| $c = 0.1, 0.2, 0.4$ | Cost of having ovulation signs present |
| $c_r = 0.4, 0.6, 0.8$ | Cost of being receptive to mating |
| $\eta = 0.1, 0.2, 0.4$ | Weighting of NGC vs. GC on a female's fitness |
| $\rho = -0.5, 0, 0.5$ | Correlation between a male's GC and NGC |
| $\hat{y} = 1$ | Mean male genetic quality |
| $\epsilon_m = 0, 0.25, 0.5$ | Parameter determining the amount of reproductive stochasticity among males |
| $\epsilon_f = 0, 0.01, 0.05$ | Parameter determining the amount of reproductive stochasticity among females |
| $\gamma = 1$ | Parameter determining the shape of the function for the amount of visual ovulation signs present |
| $\alpha = 0, 0.2, 0.6, 1$ | Parameter controlling the maximum benefit of protecting against infanticide (note $\alpha = 0$ corresponds to the case with NO infanticide) |
| $\beta = 1.25, 1.5, 1.75$ | Parameter controlling the relative maximum cost of infanticide in the paternity probability effect function |
| $\tau = 2$ | Exponent in the paternity probability effect function $g$ |
| $\omega = 1$ | Parameter in the male effectiveness function $f(j)$ |
| $\kappa = 0, 0.5, 1$ | Parameter determining the weight males put on the amount of visual ovulation signs in their paternity estimate |
| $\nu = 1$ | Migration rate (between zero and one) |
| $\mu = 0.001$ | Probability of mutation (per gene per generation) |
| $\sigma = \sqrt{0.1}$ | Standard deviation for mutation magnitude |

#### Effects of all Parameters

See Table A2 for a summary of effects for all main parameters used in the model. These effects are all illustrated for simulations with concealed ovulation fixed (Figures 2,3,4), visible ovulation signs fixed (Figures 5,6,7), and each of receptivity length, ovulation signs magnitude, and ovulation signs length evolving (Figures 8-16).

**Table A2:** Effects of All Parameters

| Traits Evolving: | Found in Figure: | Value(s) of $\rho$: | Other Parameters Investigated: |
| --- | --- | --- | --- |
| Only receptivity $r$,<br>with concealed<br>ovulation fixed<br>($m = 0, \ell = 0$) | 2 | $\rho = 0.5, 0, -0.5$ | $N, b, c_r, \eta$ |
| | 3 | $\rho = 0.5, 0, -0.5$ | $\alpha, \beta, \kappa, \eta$ |
| | 4 | $\rho = 0.5, 0, -0.5$ | $\epsilon_m, \epsilon_f, c_r, \eta$ |
| Only receptivity $r$,<br>with fixed visual<br>ovulation signs<br>($m = 0.9, \ell = 4$) | 5 | $\rho = 0.5, 0, -0.5$ | $N, b, c_r, \eta$ |
| | 6 | $\rho = 0.5, 0, -0.5$ | $\alpha, \beta, \kappa, \eta$ |
| | 7 | $\rho = 0.5, 0, -0.5$ | $\epsilon_m, \epsilon_f, c_r, \eta$ |
| Receptivity length<br>$r$, ovulation signs<br>magnitude $m$, and<br>ovulation signs<br>length $\ell$ | 8 | $\rho = 0.5$ | $N, b, c, c_r, \eta$ |
| | 9 | $\rho = 0$ | $N, b, c, c_r, \eta$ |
| | 10 | $\rho = -0.5$ | $N, b, c, c_r, \eta$ |
| | 11 | $\rho = 0.5$ | $\alpha, \beta, \kappa, \eta$ |
| | 12 | $\rho = 0$ | $\alpha, \beta, \kappa, \eta$ |
| | 13 | $\rho = -0.5$ | $\alpha, \beta, \kappa, \eta$ |
| | 14 | $\rho = 0.5$ | $\epsilon_m, \epsilon_f, c_r, \eta$ |
| | 15 | $\rho = 0$ | $\epsilon_m, \epsilon_f, c_r, \eta$ |
| | 16 | $\rho = -0.5$ | $\epsilon_m, \epsilon_f, c_r, \eta$ |

#### ANOVA Results

We ran an analysis of variance (ANOVA) to determine which of receptivity length (*r*), ovulation signs magnitude *m*, and ovulation signs length *l* are affected most by which parameters, as introduced in the main text.

The following tables (Tables A3, A4, A5) give the detailed results of these tests. Each table below reflects the results from different simulation sets. The numbers in each of these tables correspond to percentage of variance, with the sign (*±*) corresponding to the direction of the effect. Any table entry with a zero means that effect is not significant (i.e., *p >* 0.05). For example, in Table A3, 31% of the variation in receptivity length (*r*) in this simulation set can be explained by parameter *c_r_*, and another 38% by parameter *η*. Since the effect of *c_r_* is negative and *η* positive, we know that as *c_r_* increases, *r* decreases, and as *η* increases, *r* instead increases. Each of the following tables can be interpreted in this fashion.

**Table A3:** Effects of *N*, *b*, *c*, *c_r_*, *η*, *ρ* on receptivity length *R* and ovulation signs magnitude *M* and length *L*

| | $R$ | $M$ | $L$ |
| --- | --- | --- | --- |
| $N$ | 0 | 0.019 | 0 |
| $b$ | 0 | 0.261 | 0.179 |
| $c$ | 0 | -0.171 | -0.313 |
| $c_r$ | -0.313 | -0.072 | -0.124 |
| $\eta$ | 0.380 | 0.005 | 0.105 |
| $\rho$ | 0 | 0.014 | 0.029 |
| error | 0.306 | 0.459 | 0.530 |

**Table A4:** Effects of *α*, *β*, *κ*, *η*, *ρ* on receptivity length *R* and ovulation signs magnitude *M* and length *L*

| | $R$ | $M$ | $L$ |
| --- | --- | --- | --- |
| $\alpha$ | 0.025 | 0.326 | 0.187 |
| $\beta$ | 0 | 0.021 | 0.007 |
| $\kappa$ | -0.132 | 0.422 | -0.294 |
| $\eta$ | 0.634 | -0.022 | 0.270 |
| $\rho$ | 0.0002 | 0 | 0.020 |
| error | 0.207 | 0.209 | 0.222 |

**Table A5:** Effects of *∊_m_*, *∊_f_*, *c_r_*, *η*, *ρ* on receptivity length *R* and ovulation signs magnitude *M* and length *L*

| | $R$ | $M$ | $L$ |
| --- | --- | --- | --- |
| $\epsilon_m$ | 0 | -0.148 | -0.017 |
| $\epsilon_f$ | 0 | -0.011 | -0.118 |
| $c_r$ | -0.317 | -0.151 | -0.122 |
| $\eta$ | 0.375 | 0 | 0.117 |
| $\rho$ | 0 | 0.021 | 0.034 |
| error | 0.308 | 0.661 | 0.593 |

## Supplementary Figures

**Figure 2:**
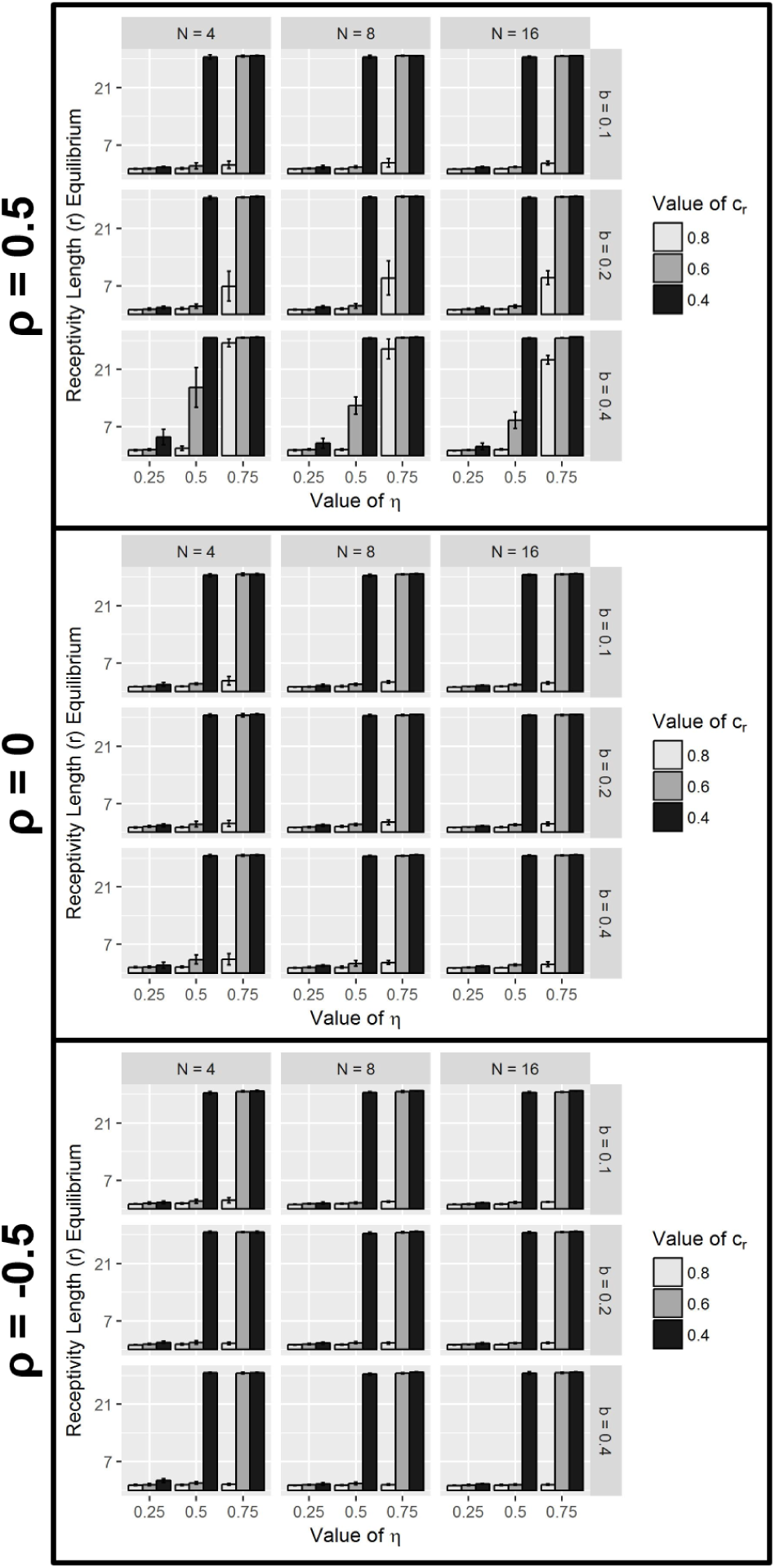
The effects of parameters *N* (group size), *b* (benefit of male genetic quality), *c_r_* (cost of receptivity length), and *η* (relative weighting of NGC) on the average equilibria values of receptivity length (*r*) for three different values of *ρ* (correlation between males’ GC and NGC), when concealed ovulation is fixed (*m* = 0, *l* = 0) for all females. Equilibria are obtained by averaging over 10 initial condition runs (with standard deviation indicated by error bars). All other parameters were held constant: *α* = 0, *c* = 0.2, *∊_m_* = 0.25, *∊_f_* = 0.01.

**Figure 3:**
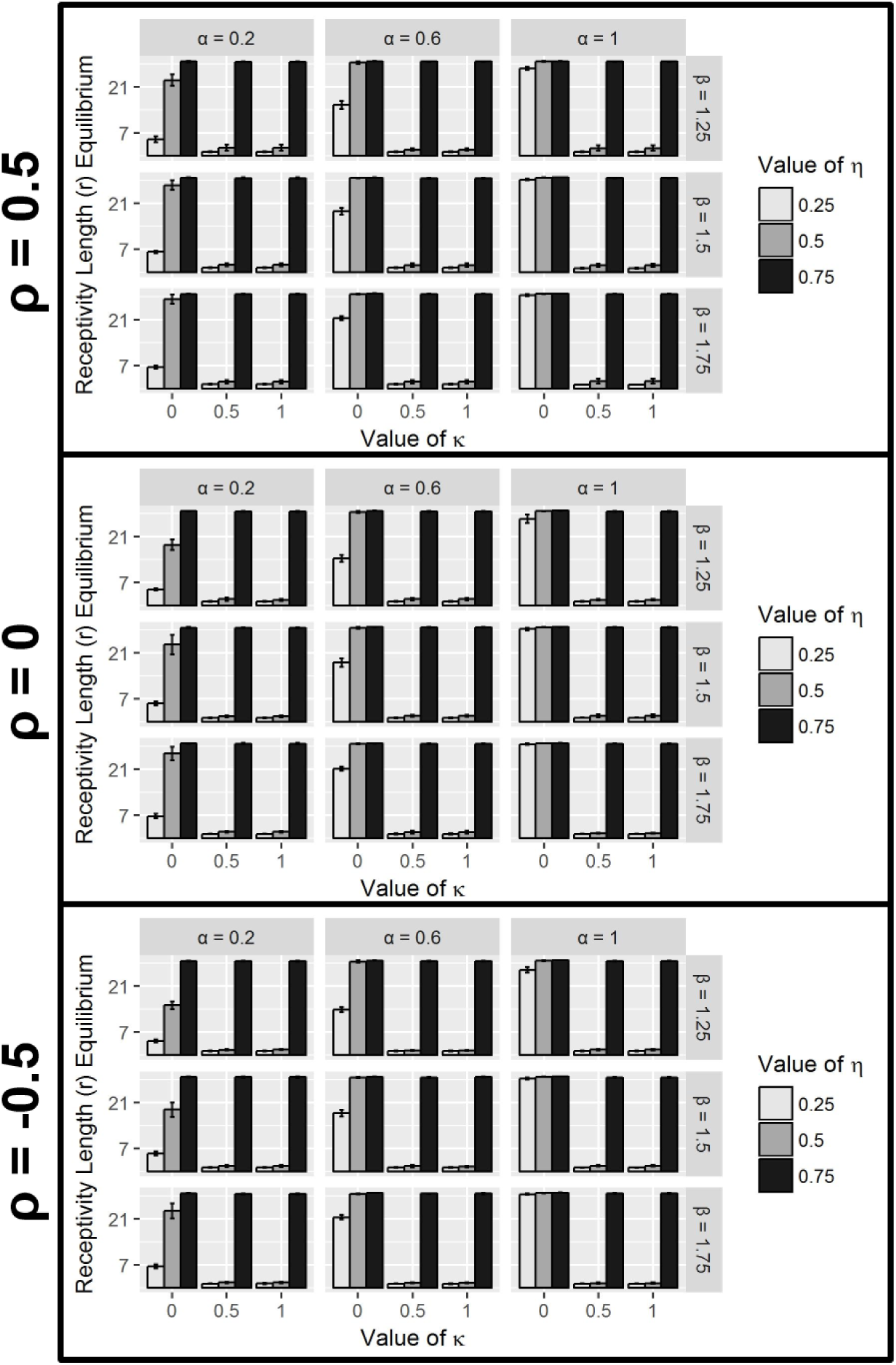
The effects of parameters *α* (maximum benefit of protecting offspring from infanticide), *β* (maximum cost of not protecting offspring from infanticide), *κ* (weight males put on visible ovulation signs when estimating their paternity), and *η* (relative weighting of NGC) on the average equilibria values of receptivity length (*r*) for three different values of *ρ* (correlation between males’ GC and NGC), when concealed ovulation is fixed (*m* = 0, *l* = 0) for all females. Equilibria are obtained by averaging over 10 initial condition runs (with standard deviation indicated by error bars). All other parameters were held constant: *N* = 8, *b* = 0.2, *c* = 0.2, *c_r_* = 0.6, *∊_m_* = 0.25, *∊_f_* = 0.01.

**Figure 4:**
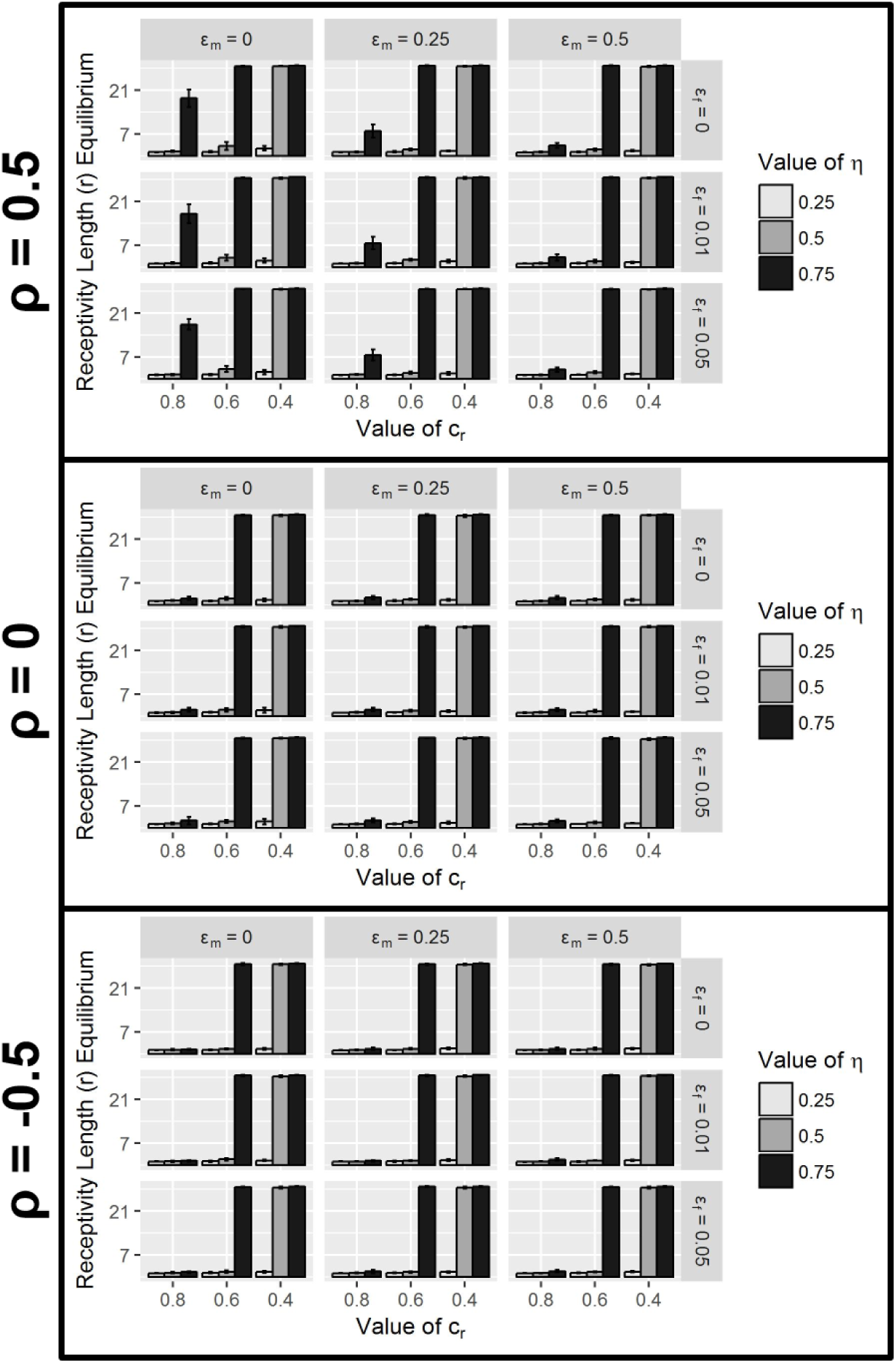
The effects of parameters *∊_m_* (male reproductive stochasticity), *∊_f_* (female reproductive stochasticity), *c_r_* (cost of receptivity length), and *η* (relative weighting of NGC) on the average equilibria values of receptivity length (*r*) for three different values of *ρ* (correlation between males’ GC and NGC), when concealed ovulation is fixed (*m* = 0, *l* = 0) for all females. Equilibria are obtained by averaging over 10 initial condition runs (with standard deviation indicated by error bars). All other parameters were held constant: *α* = 0, *N* = 8, *b* = 0.2, *c* = 0.2.

**Figure 5:**
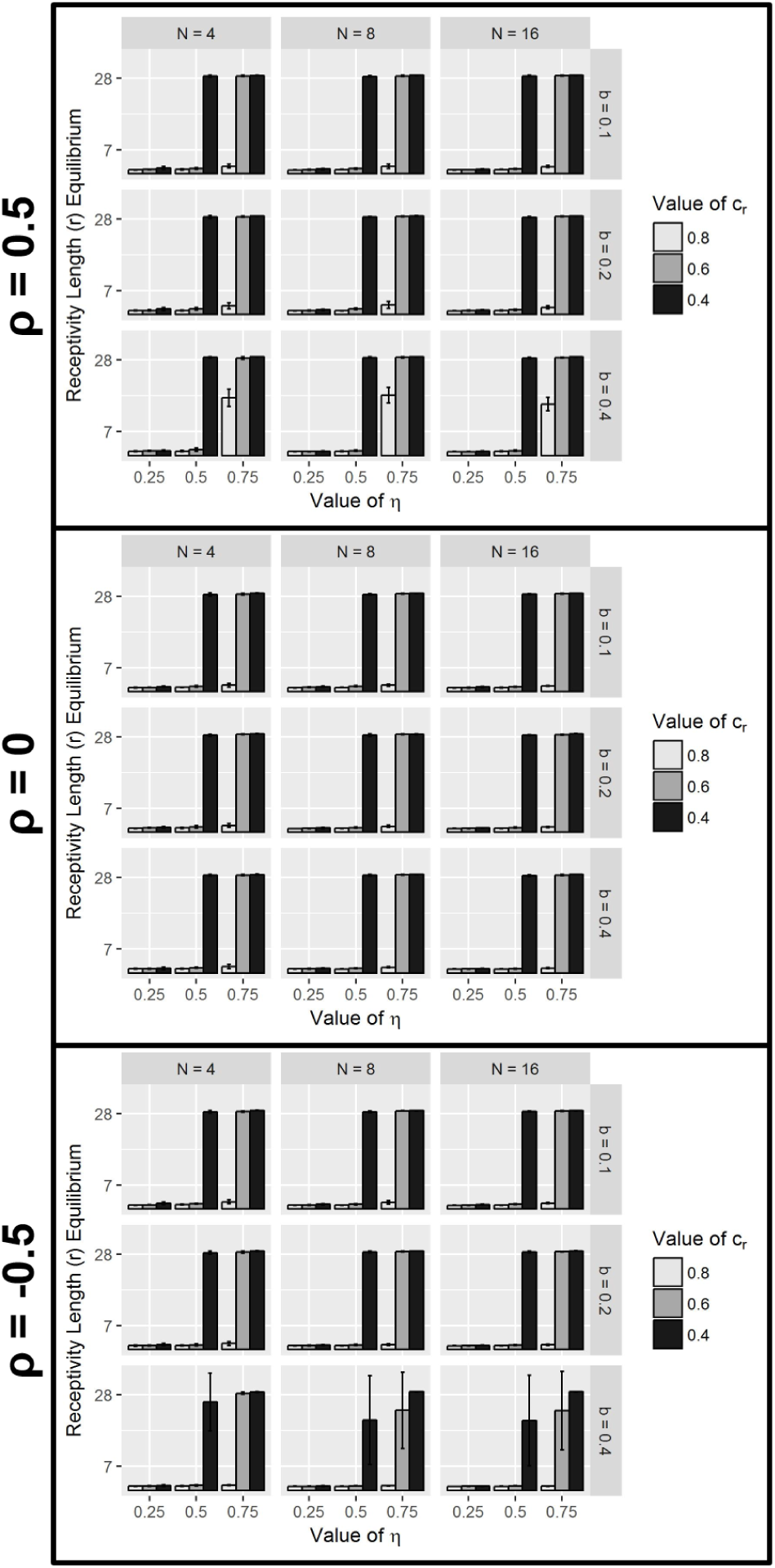
The effects of parameters *N* (group size), *b* (benefit of male genetic quality), *c_r_* (cost of receptivity length), and *η* (relative weighting of NGC) on the average equilibria values of receptivity length (*r*) for three different values of *ρ* (correlation between males’ GC and NGC), when visible ovulation signs are fixed (*m* = 0.9, *l* = 4) for all females. Equilibria are obtained by averaging over 10 initial condition runs (with standard deviation indicated by error bars). All other parameters were held constant: *α* = 0, *c* = 0.2, *∊_m_* = 0.25, *∊_f_* = 0.01.

**Figure 6:**
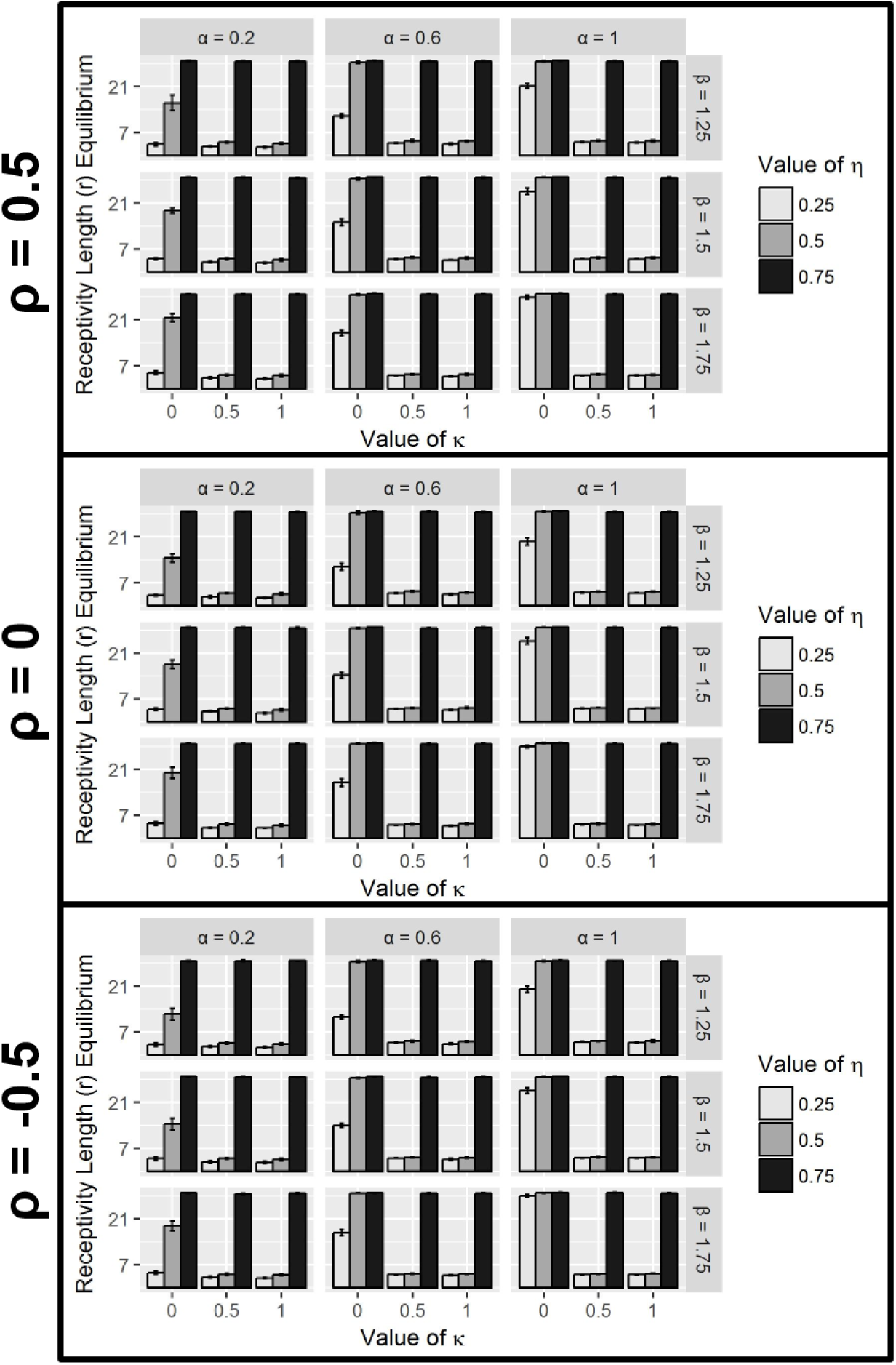
The effects of parameters *α* (maximum benefit of protecting offspring from infanticide), *β* (maximum cost of not protecting offspring from infanticide), *κ* (weight males put on visible ovulation signs when estimating their paternity), and *η* (relative weighting of NGC) on the average equilibria values of receptivity length (*r*) for three different values of *ρ* (correlation between males’ GC and NGC), when visible ovulation signs are fixed (*m* = 0.9, *l* = 4) for all females. Equilibria are obtained by averaging over 10 initial condition runs (with standard deviation indicated by error bars). All other parameters were held constant: *N* = 8, *b* = 0.2, *c* = 0.2, *c_r_* = 0.6, *∊_m_* = 0.25, *∊_f_* = 0.01.

**Figure 7:**
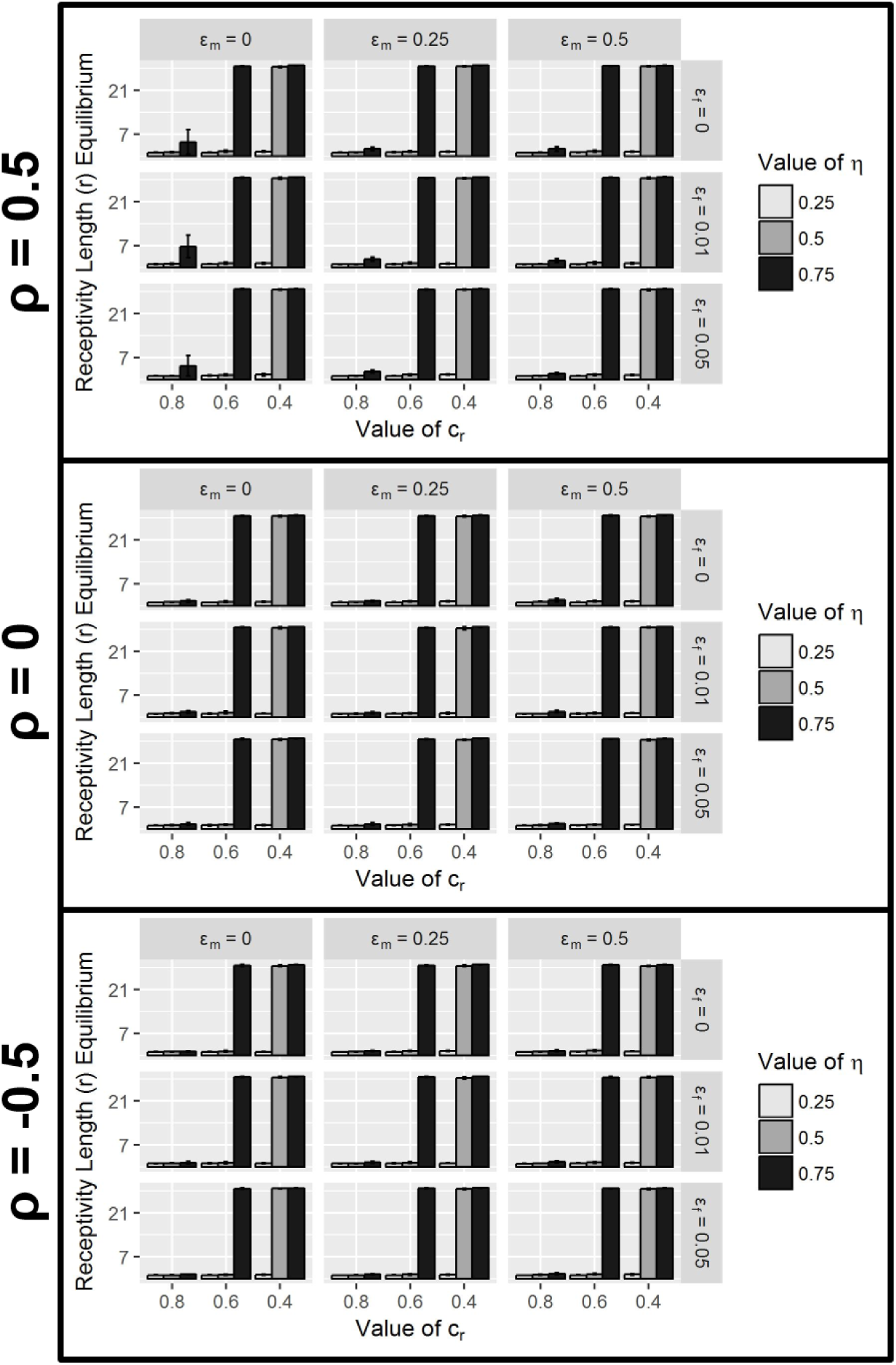
The effects of parameters *∊_m_* (male reproductive stochasticity), *∊_f_* (female reproductive stochasticity), *c_r_* (cost of receptivity length), and *η* (relative weighting of NGC) on the average equilibria values of receptivity length (*r*) for three different values of *ρ* (correlation between males’ GC and NGC), when visible ovulation signs are fixed (*m* = 0.9, *l* = 4) for all females. Equilibria are obtained by averaging over 10 initial condition runs (with standard deviation indicated by error bars). All other parameters were held constant: *α* = 0, *N* = 8, *b* = 0.2, *c* = 0.2.

**Figure 8:**
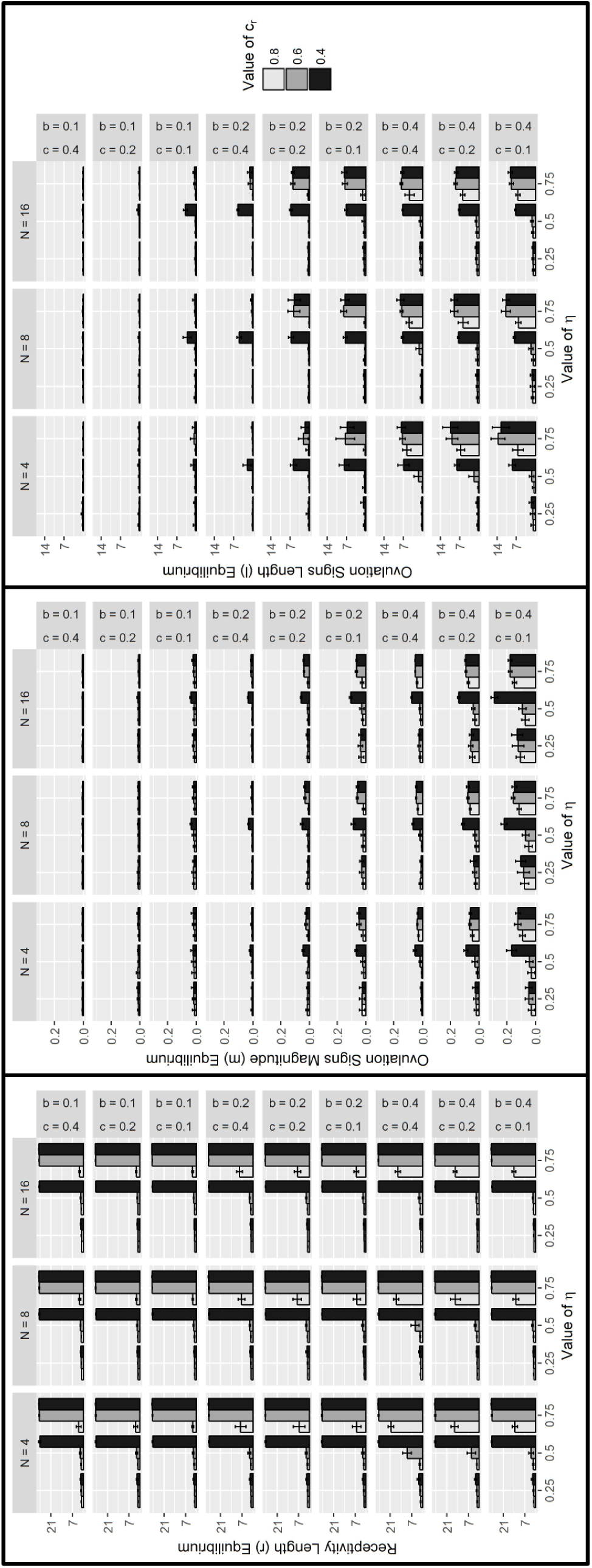
The effects of parameters *N* (group size), *b* (benefit of male genetic quality), *c* (cost of visible ovulation signs), *c_r_* (cost of receptivity length), and *η* (relative weighting of NGC) on the average equilibria values of receptivity length (*r*), ovulation signs magnitude (*m*), and ovulation signs length (*l*) with *ρ* = 0.5 (correlation between males’ GC and NGC). Equilibria are obtained by averaging over 16 initial condition runs (with standard deviation indicated by error bars). All other parameters were held constant: *α* = 0, *∊_m_* = 0.25, *∊_f_* = 0.01.

**Figure 9:**
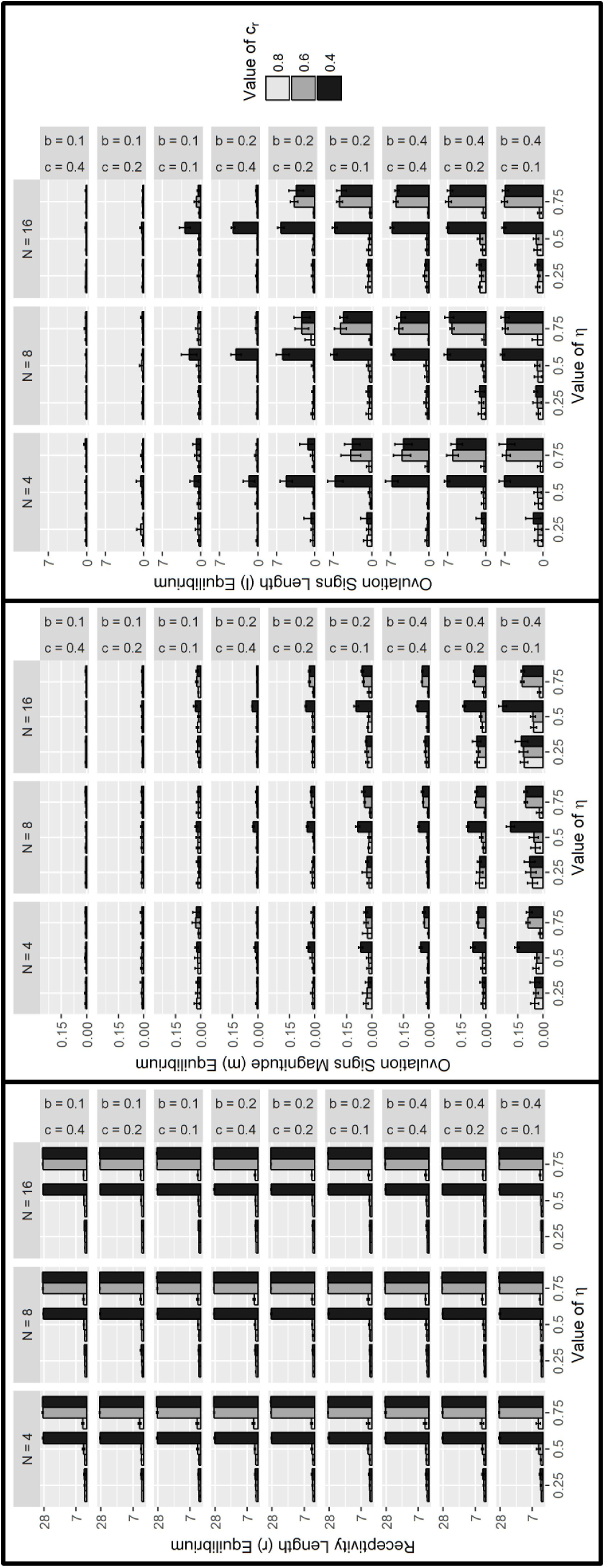
The effects of parameters *N* (group size), *b* (benefit of male genetic quality), *c* (cost of visible ovulation signs), *c_r_* (cost of receptivity length), and *η* (relative weighting of NGC) on the average equilibria values of receptivity length (*r*), ovulation signs magnitude (*m*), and ovulation signs length (*l*) with *ρ* = 0 (correlation between males’ GC and NGC). Equilibria are obtained by averaging over 16 initial condition runs (with standard deviation indicated by error bars). All other parameters were held constant: *α* = 0, *∊_m_* = 0.25, *∊_f_* = 0.01.

**Figure 10:**
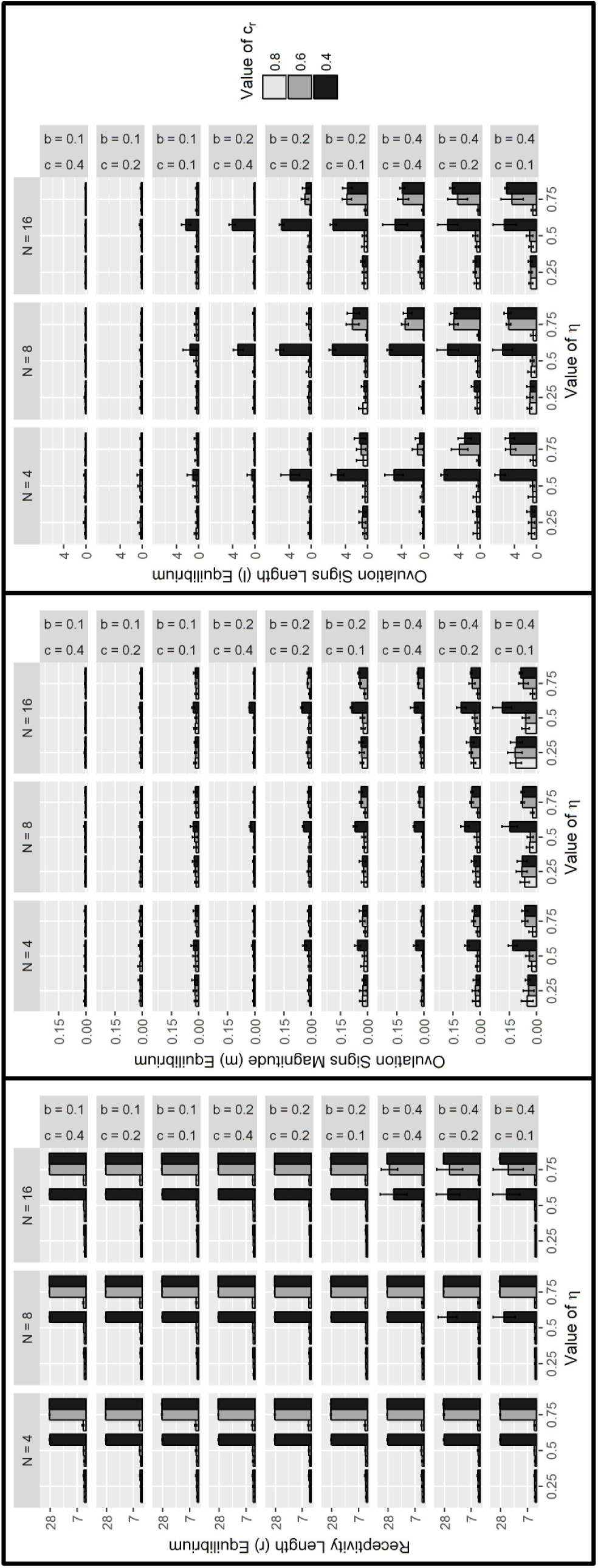
The effects of parameters *N* (group size), *b* (benefit of male genetic quality), *c* (cost of visible ovulation signs), *c_r_* (cost of receptivity length), and *η* (relative weighting of NGC) on the average equilibria values of receptivity length (*r*), ovulation signs magnitude (*m*), and ovulation signs length (*l*) with *ρ* = −0.5 (correlation between males’ GC and NGC). Equilibria are obtained by averaging over 16 initial condition runs (with standard deviation indicated by error bars). All other parameters were held constant: *α* = 0, *∊_m_* = 0.25, *∊_f_* = 0.01.

**Figure 11:**
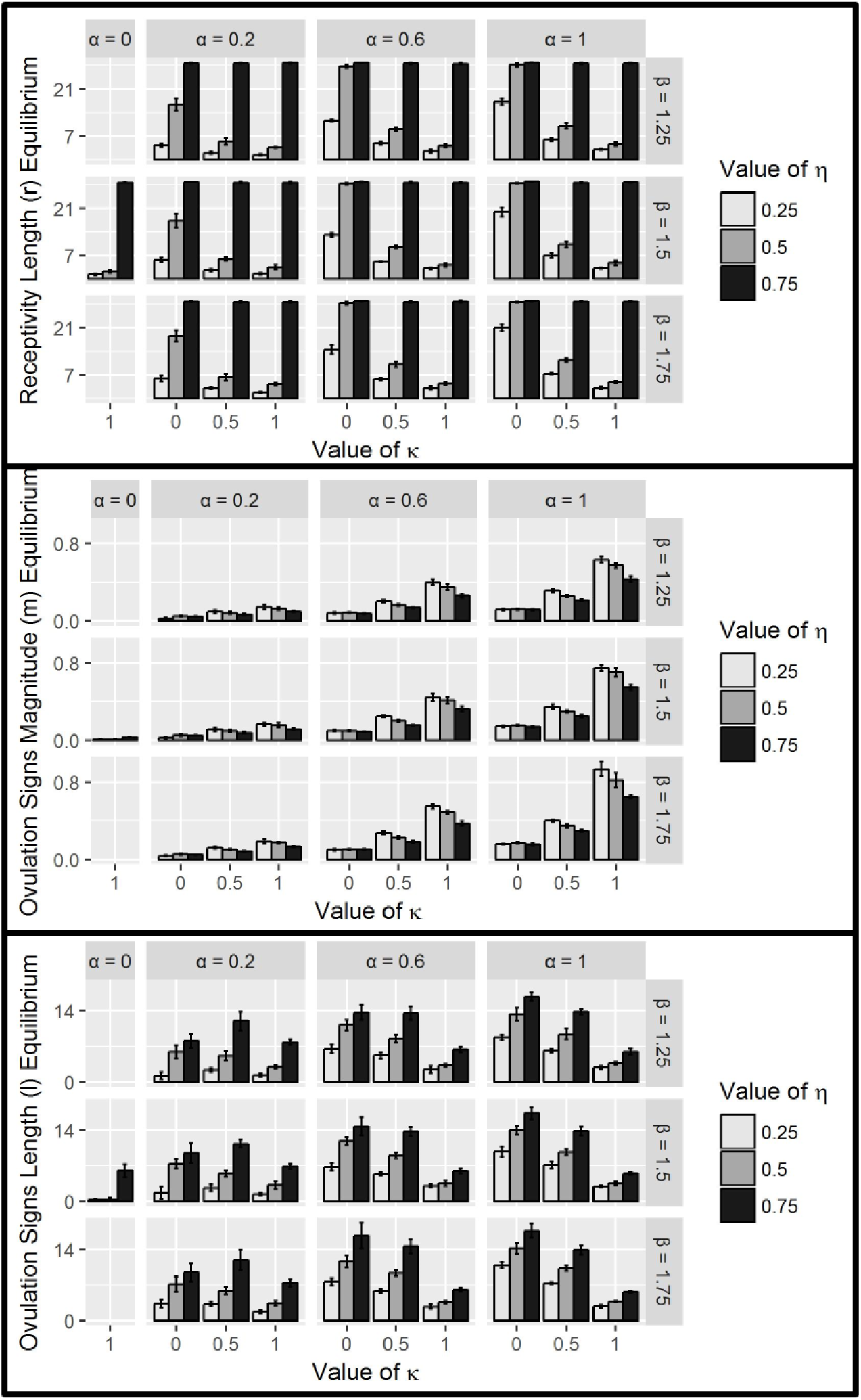
The effects of parameters *α* (maximum benefit of infanticide), *β* (proportional to the maximum cost of infanticide), *κ* (weight males put on females having ovulation signs visible), and *η* (relative weighting of NGC) on the average equilibria values of receptivity length (*r*), ovulation signs magnitude (*m*), and ovulation signs length (*l*) with *ρ* = 0.5 (correlation between males’ GC and NGC). Equilibria are obtained by averaging over 16 initial condition runs (with standard deviation indicated by error bars). All other parameters were held constant: *N* = 8, *b* = 0.2, *c* = 0.2, *c_r_* = 0.6, *∊_m_* = 0.25, *∊_f_* = 0.01.

**Figure 12:**
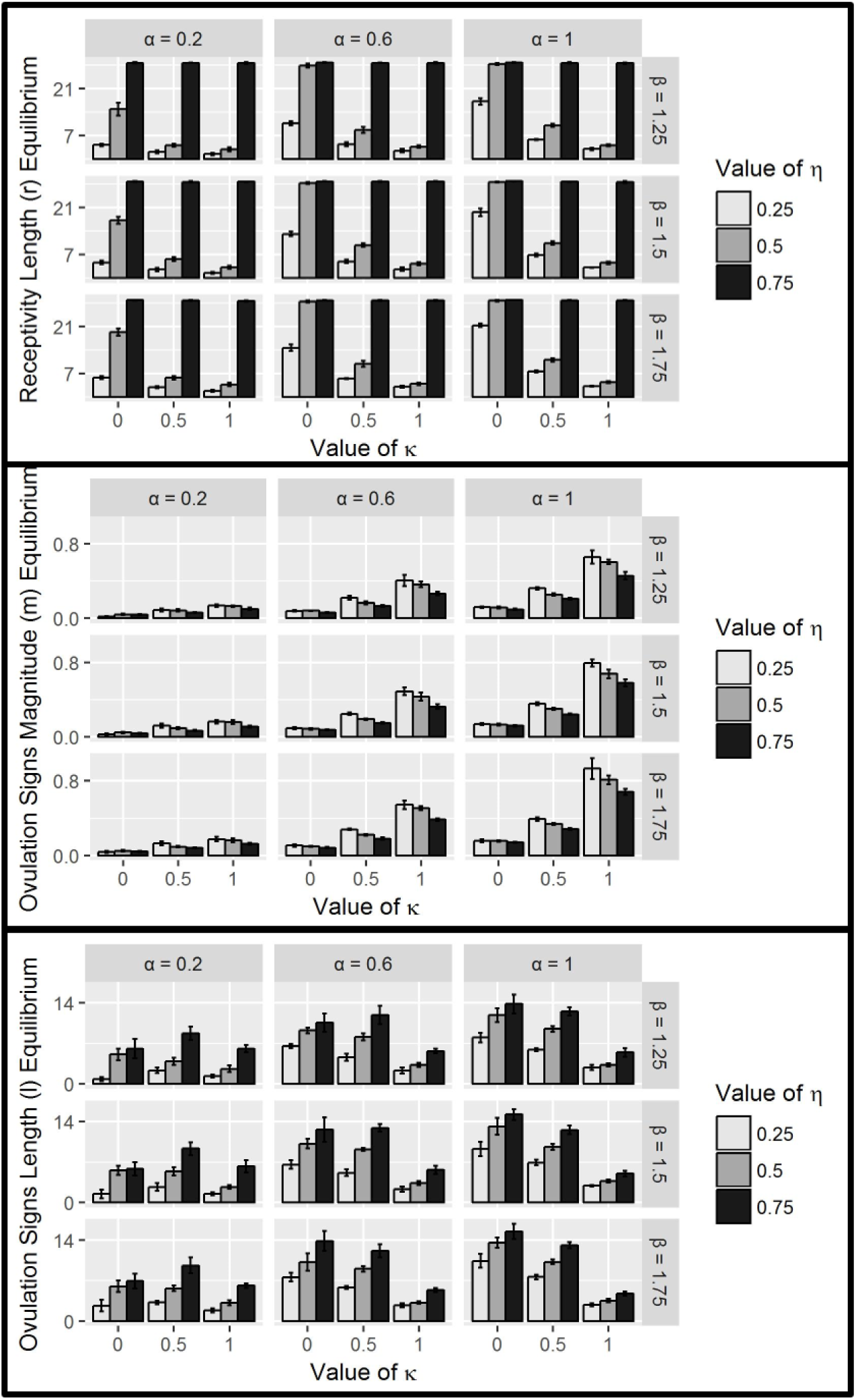
The effects of parameters *α* (maximum benefit of infanticide), *β* (proportional to the maximum cost of infanticide), *κ* (weight males put on females having ovulation signs visible), and *η* (relative weighting of NGC) on the average equilibria values of receptivity length (*r*), ovulation signs magnitude (*m*), and ovulation signs length (*l*) with *ρ* = 0 (correlation between males’ GC and NGC). Equilibria are obtained by averaging over 16 initial condition runs (with standard deviation indicated by error bars). All other parameters were held constant: *N* = 8, *b* = 0.2, *c* = 0.2, *c_r_* = 0.6, *∊_m_* = 0.25, *∊_f_* = 0.01.

**Figure 13:**
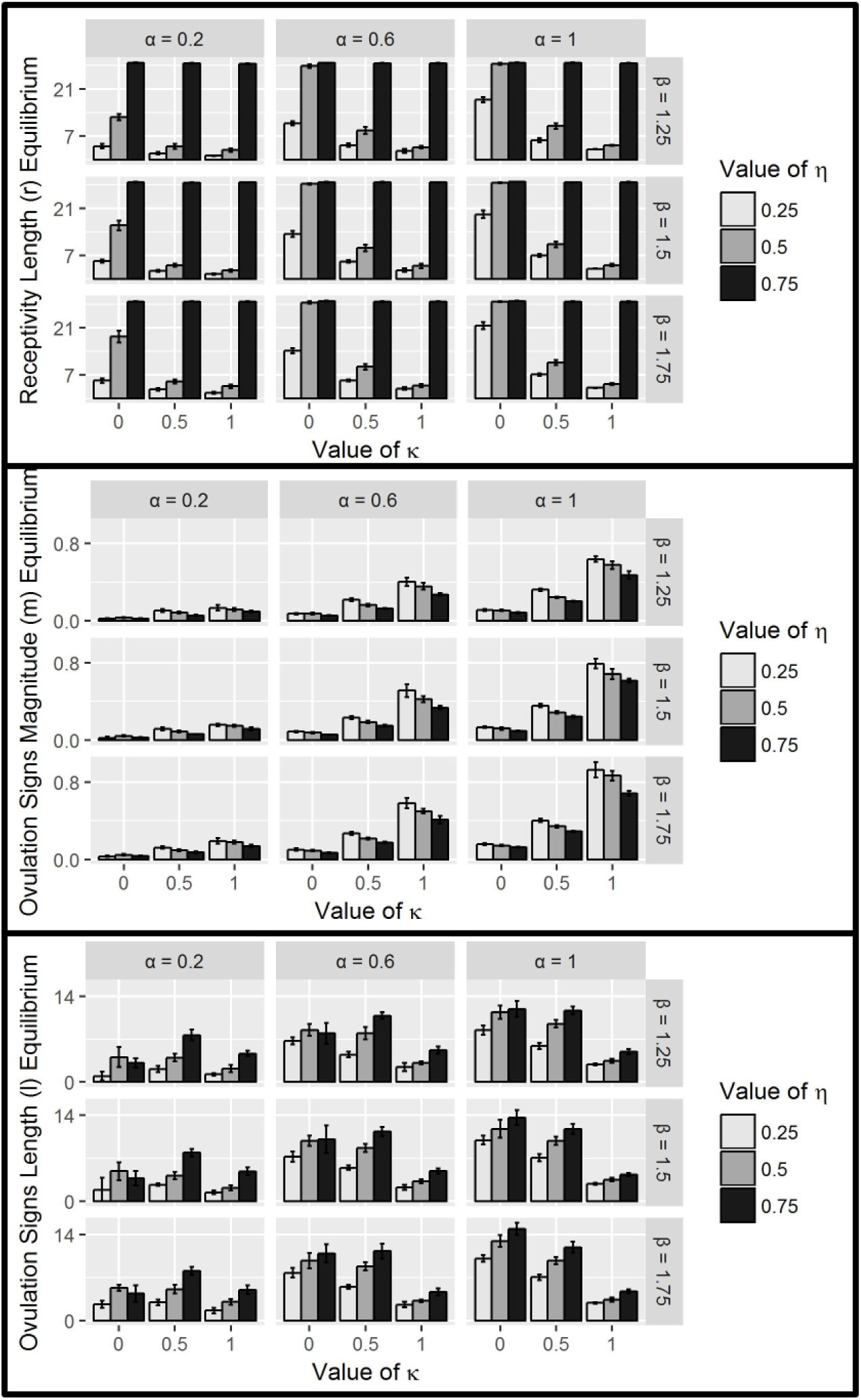
The effects of parameters *α* (maximum benefit of infanticide), *β* (proportional to the maximum cost of infanticide), *κ* (weight males put on females having ovulation signs visible), and *η* (relative weighting of NGC) on the average equilibria values of receptivity length (*r*), ovulation signs magnitude (*m*), and ovulation signs length (*l*) with *ρ* = −0.5 (correlation between males’ GC and NGC). Equilibria are obtained by averaging over 16 initial condition runs (with standard deviation indicated by error bars). All other parameters were held constant: *N* = 8, *b* = 0.2, *c* = 0.2, *c_r_* = 0.6, *∊_m_* = 0.25, *∊_f_* = 0.01.

**Figure 14:**
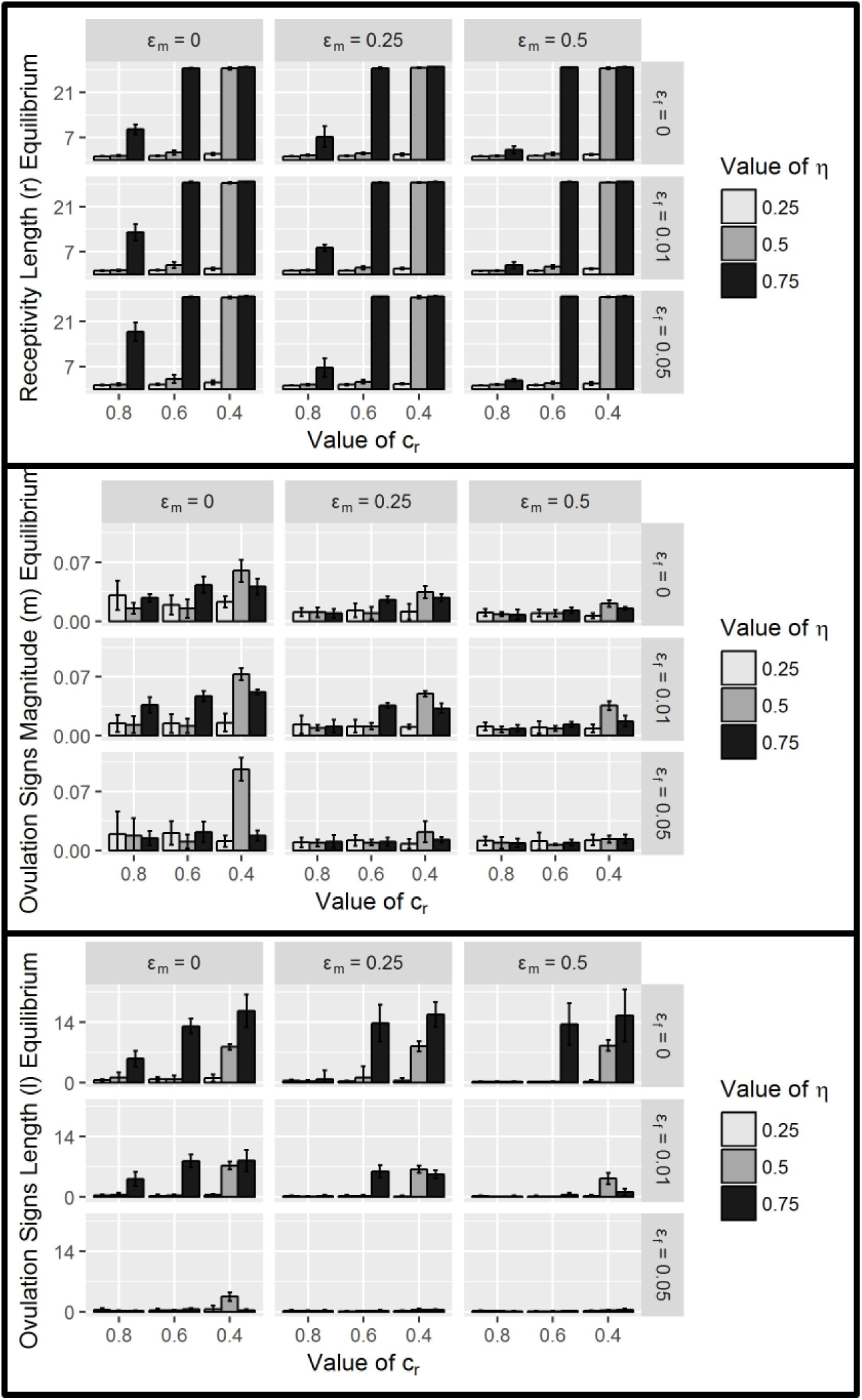
The effects of parameters *∊_m_* (reproductive stochasticity among males), *∊_f_* (reproductive stochasticity among females), *c_r_* (cost of receptivity length), and *η* (relative weighting of NGC) on the average equilibria values of receptivity length (*r*), ovulation signs magnitude (*m*), and ovulation signs length (*l*) with *ρ* = 0.5 (correlation between males’ GC and NGC). Equilibria are obtained by averaging over 16 initial condition runs (with standard deviation indicated by error bars). All other parameters were held constant: *N* = 8, *b* = 0.2, *c* = 0.2, *α* = 0.

**Figure 15:**
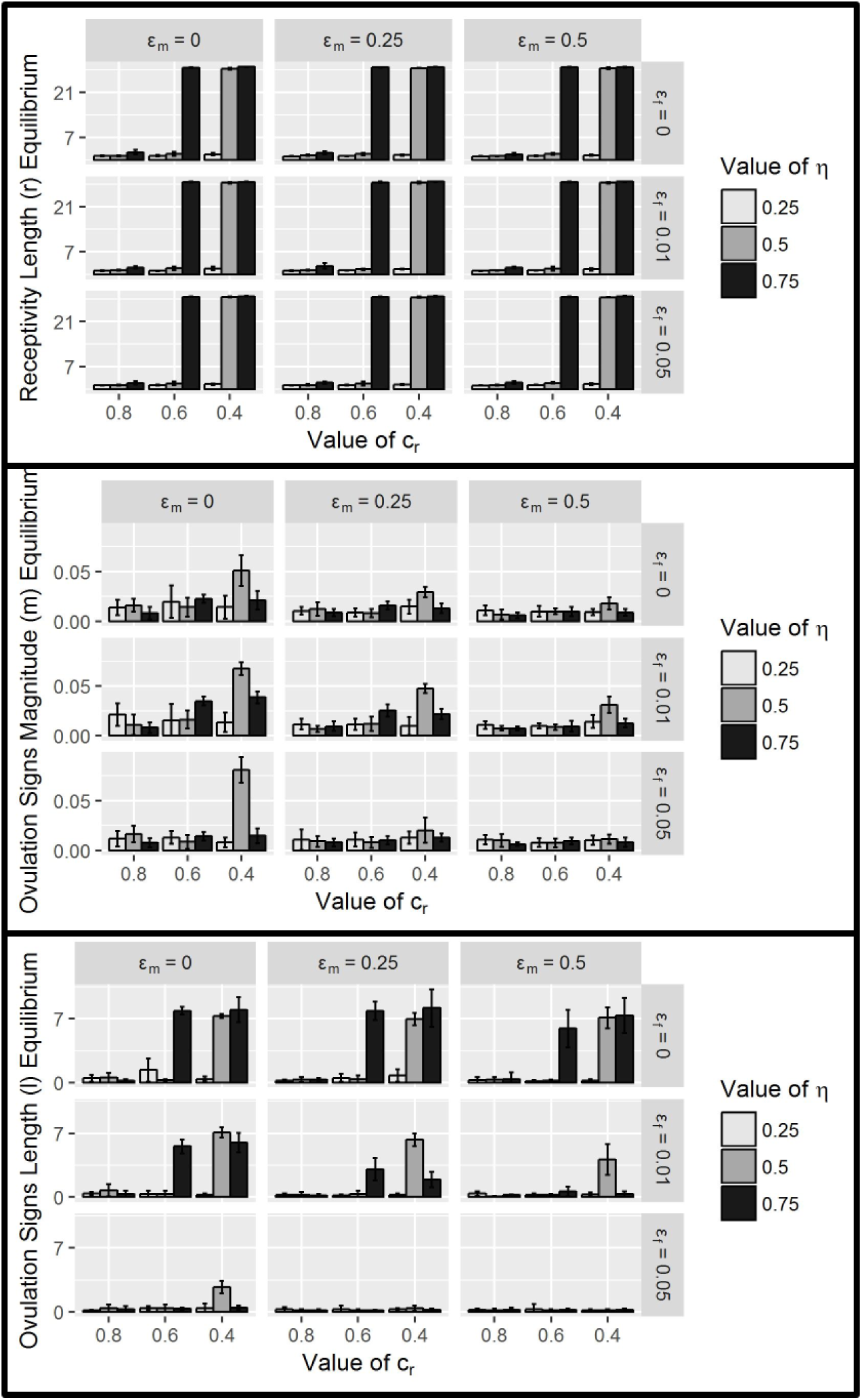
The effects of parameters *∊_m_* (reproductive stochasticity among males), *∊_f_* (reproductive stochasticity among females), *c_r_* (cost of receptivity length), and *η* (relative weighting of NGC) on the average equilibria values of receptivity length (*r*), ovulation signs magnitude (*m*), and ovulation signs length (*l*) with *ρ* = 0 (correlation between males’ GC and NGC). Equilibria are obtained by averaging over 16 initial condition runs (with standard deviation indicated by error bars). All other parameters were held constant: *N* = 8, *b* = 0.2, *c* = 0.2, *α* = 0.

**Figure 16:**
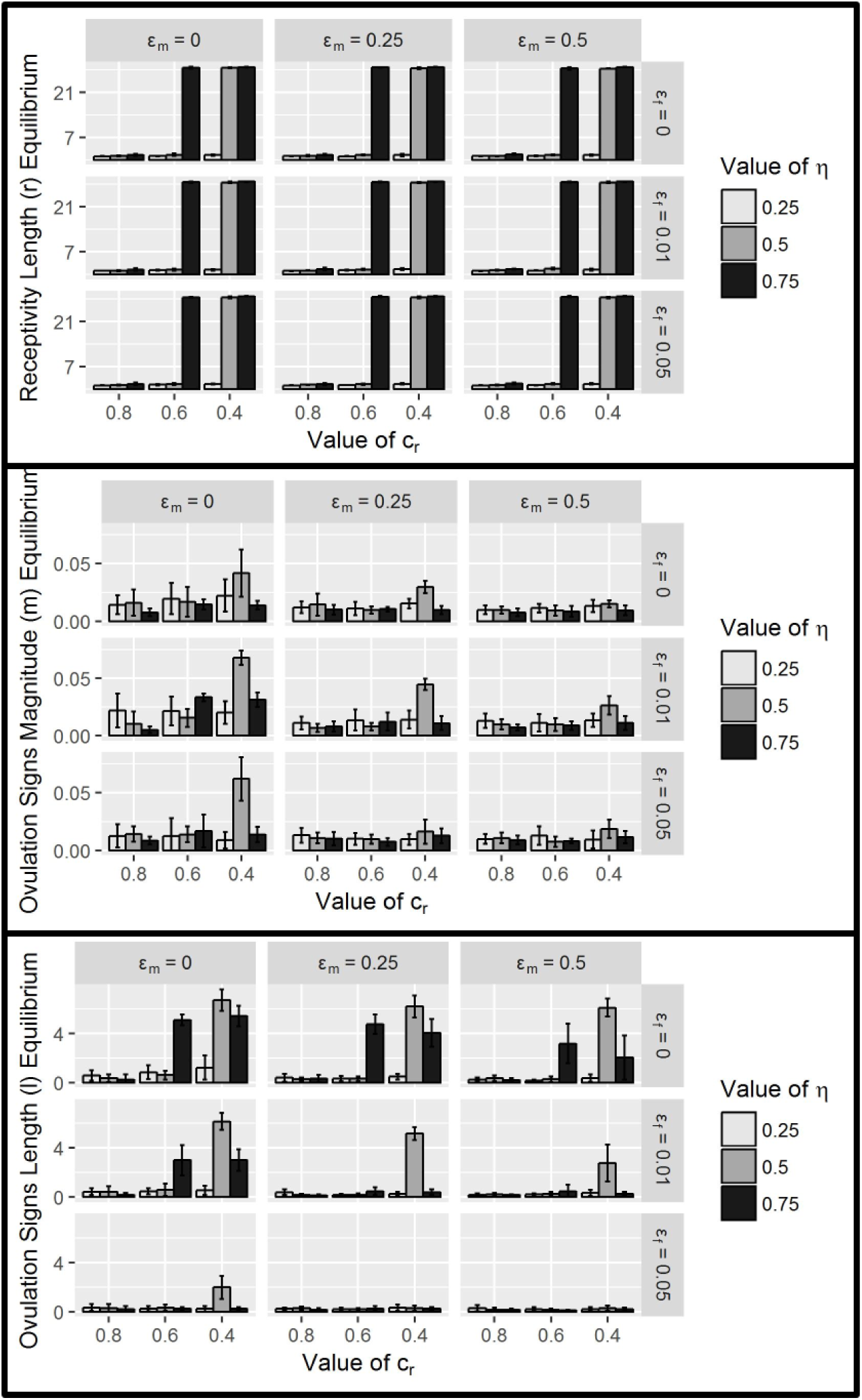
The effects of parameters *∊_m_* (reproductive stochasticity among males), *∊_f_* (reproductive stochasticity among females), *c_r_* (cost of receptivity length), and *η* (relative weighting of NGC) on the average equilibria values of receptivity length (*r*), ovulation signs magnitude (*m*), and ovulation signs length (*l*) with *ρ* = −0.5 (correlation between males’ GC and NGC). Equilibria are obtained by averaging over 16 initial condition runs (with standard deviation indicated by error bars). All other parameters were held constant: *N* = 8, *b* = 0.2, *c* = 0.2, *α* = 0.

